# Matchtigs: minimum plain text representation of kmer sets

**DOI:** 10.1101/2021.12.15.472871

**Authors:** Sebastian Schmidt, Shahbaz Khan, Jarno Alanko, Giulio E. Pibiri, Alexandru I. Tomescu

## Abstract

We propose a polynomial algorithm computing a *minimum* plain-text representation of kmer sets, as well as an efficient near-minimum greedy heuristic. When compressing read sets of large model organisms or bacterial pangenomes, with only a minor runtime increase, we shrink the representation by up to 60% over unitigs and 27% over previous work. Additionally, the number of strings is decreased by up to 97% over unitigs and 91% over previous work. Finally, a small representation has advantages in downstream applications, as it speeds up SSHash-Lite queries by up to 4.26× over unitigs and 2.10× over previous work.

**Availability:** matchtigs: https://github.com/algbio/matchtigs

SSHash-Lite: https://github.com/jermp/sshash-lite

## 1 Background

### Motivation

The field of kmer-based methods has seen a surge of publications in the last years. Examples include alignment-free sequence comparison [1, 2, 3], variant calling and genotyping [4, 5, 6, 7, 8], transcript abundance estimation [9], metagenomic classification [10, 11, 12, 13], abundance profile inference [14], indexing of variation graphs [15, 16], estimating the similarity between metagenomic datasets [17], species identification [18, 19] and sequence alignment to de Bruijn graphs [20, 21, 22, 23]. All these methods are based mainly on kmer sets, i.e. on the existence or non-existence of kmers. They ignore further information like for example predecessor and successor relations between kmers which are represented by the topology of a de Bruijn graph [24, 25, 26].

On the other hand, many classical methods such as genome assemblers [26, 27, 28, 29, 30, 31, 32, 33, 34, 35] and related algorithms [36, 37, 38], are based on de Bruijn graphs and their topology. To increase the efficiency of these methods, the graphs are usually compacted by contracting all paths where all inner nodes have in- and outdegree one. These paths are commonly known as *unitigs*, and their first usage can be traced back to [39]. Since unitigs contain no branches in their inner nodes, they do not alter the topology of the graph, and in turn enable the exact same set of analyses. There are highly engineered solutions available to compute a compacted de Bruijn graph by computing unitigs from any set of strings in memory [23] or with external memory [33, 40, 41]. Incidentally, the set of unitigs computed from a set of strings is also a way to store a set of kmers without repetition, and thus in reasonably small space. However, the necessity to preserve the topology of the graph makes unitigs an inferior choice to represent kmer sets, as the sum of their length is still far from optimal, and they consist of many separate strings. The possibility to ignore the topology for kmer-based methods opens more leeway in their representation that can be exploited to reduce the resource consumption of existing and future bioinformatics tools.

The need for such representations becomes apparent when observing the amount of data available to bioinformaticians. For example, the number of complete bacterial genomes available in RefSeq [42] more than doubled between May 2020 and July 2021 from around 9000 [43] to around 21000^[1]^. And with the ready availability of modern sequencing technologies, the amount of genomic data will increase further in the next years. In turn, analysing this data requires an ever growing amount of computational resources. But this could be relieved through a smaller representation that reduces the RAM usage and speeds up the analysis tools, and thereby allows to run larger pipelines using less computational resources. To fulfil this goal, a *plain text* representation would be the most useful: if the representation has to be decompressed before usage, then this likely erases the savings in RAM and/or adds additional runtime overhead. Formally, a plain text representation is a set of strings that contains each kmer from the input strings (forward, reverse-complemented, or both) and no other kmer. We denote such a set as a *spectrum preserving string set* (SPSS), borrowing the naming from [44] and redefining it slightly^[2]^. Such a plain text representation has the great advantage that some tools (like e.g. Bifrost’s query [23]) can use it without modification. We expect that even those tools that require modifications would not be hard to modify (like e.g. SSHash [45] which we modified here as an example).

### Related work

The concept of storing a set of kmers in plain text without repeating kmers to achieve a smaller and possibly simpler representation has recently been simultaneously discovered and named *spectrum preserving string sets [without kmer repetition]* by Rahman and Medvedev [44] as well as *simplitigs* by Břinda, Baym and Kucherov [43]. To avoid confusion with our redefinition of the SPSS, we call this concept *simplitigs* in our work. Both Rahman and Medvedev and Břinda, Baym and Kucherov propose an implementation that greedily joins consecutive unitigs to compute such a representation. The UST algorithm by Rahman and Medvedev works on the node-centric de Bruijn graph of the input strings and finds arbitrary paths in the graph starting from arbitrary nodes. Each node is visited exactly by one path, and whenever a path cannot be extended forwards (because a dead-end was found, or all successor nodes have been visited already), then a new path is started from a new random node. Before a new path is started this way, if any successor node of the finished path marks the start of a different path, then the two paths are joined. During the traversal, the unitigs of the visited nodes are concatenated (without repeating the *k —* 1 overlapping characters) and those strings are the final output. Brinda, Baym and Kucherov’s greedy algorithm to compute simplitigs (for which the authors provide an implementation under the name ProphAsm [43]) does not construct a de Bruijn graph, but instead collects all kmers into a hash table. Then it extends arbitrary kmers both forwards and backwards arbitrarily until they cannot be extended anymore, without repeating any kmers. The extended kmers are the final output.

Both heuristics greatly reduce the number of strings *(string count*, SC) as well as the total amount of characters in the strings *(cumulative length*, CL) required to store a kmer set. The reduction in CL directly relates to a lower memory consumption for storing a set of strings, but also the reduction in SC is very useful. When storing a set of strings, not only the strings need to be stored, but also some index structure telling where they start and end. This structure can be smaller if less strings exist^[3]^. Břinda, Baym and Kucherov show that both SC and CL are greatly reduced for very tangled de Bruijn graphs, like graphs for single large genomes with small kmer length and pangenome graphs with many genomes. Additionally they show merits of using heuristic simplitigs in downstream applications like an improvement in run time of kmer queries using BWA [46], as well as a reduction in space required when storing heuristic simplitigs compressed with general-purpose compressors over storing unitigs compressed in the same way. Rahman and Medvedev show a significant reduction in SC and CL on various data sets as well, and also show a reduction in space required to store heuristic simplitigs over unitigs when compressed with general-purpose compressors.

The authors of both papers also give a lower bound on the cumulative length of simplitigs, and show that their heuristics achieve representations with a cumulative length very close to the lower bound for typical values of *k* (31 for bacterial genomes and 51 for eukaryotic genomes). Brinda, Baym and Kucherov also experiment with lower values of *k* (< 20 for bacterial genomes and < 30 for eukaryotic genomes) which make the de Bruijn graph more dense to almost complete, and show that in these cases, their heuristic does not get as close to the lower bound as for larger values of *k*. Further, the authors of both papers consider whether computing minimum simplitigs without repeating kmers might be NP-hard. This has recently been disproven by Schmidt and Alanko [47], and in fact simplitigs with minimum cumulative length can be computed in linear time^[4]^. Their algorithm constructs a bidirected arc-centric de Bruijn graph in linear time using a suffix tree, and then Eulerises it by inserting *breaking arcs*. It then computes a bidirected Eulerian circuit in the Eulerised graph and breaks it at all breaking arcs. The strings spelled by the resulting walks are the optimal simplitigs, named *Eulertigs*. Specifically, they leave out all parts from the matchtigs algorithm that relate to concatenating unitigs by repeating kmers, and instead only concatenate consecutive unitigs in an optimal way. In line with the previous results about the lower bounds [44, 43], Eulertigs are only marginally smaller than the strings computed by previous heuristics. All these suggest that no further progress is possible when kmer repetitions are not allowed in a plain text representation.

There are already tools available that use simplitigs. The compacted de Bruijn graph builder cuttlefish2 [40] has an option to output simplitigs instead of maximal unitigs. A recent proposal for a standardised file format for kmer sets explicitly supports simplitigs [48]. Also the kmer dictionary SSHash [45] uses simplitigs to achieve a smaller representation and to answer queries more efficiently. Here, the higher efficiency is achieved both by reducing the space required to store the kmers themselves, but also due to the lower string count reducing the size of the index data structures on top. Further, a recent proposal to index genomic sequences as opposed to kmer sets works with simplitigs without modification [49], and with minor extra book-keeping also for general SPSSs. In that work, the size of the SPSS is very minor compared to the size of the index, however, major components of the index may be smaller if the SPSS contains less strings, which can be achieved by using greedy matchtigs. Our algorithms were also integrated into the external-memory de Bruijn graph compactor GGCAT [41], which was easy to do^[5]^.

In the wider field of finding small representations of kmer sets that are not necessarily in plain text, there exists for example ESSCompress [50], which uses an extended DNA alphabet to encode similar kmers in smaller space. Another non-plain text representation is REINDEER [51], which uses substrings of unitigs with kmers of similar abundance to not just store the existence of kmers, but also their abundance in each single genome of a pangenome. Lastly, in [52] the authors use an algorithm similar to ProphAsm and UST to compress multiple kmer sets by separating the unique kmer content of each set from the kmer content shared with other sets.

### Our contribution

In this paper we propose the first algorithm to find an SPSS of *minimum* size (CL). Moreover, we show that a minimum SPSS with repeated kmers is polynomially solvable, based on a many-to-many min-cost path query and a min-cost perfect matching approach. We further propose a faster and more memory-efficient greedy heuristic to compute a small SPSS that skips the optimal matching step, but still produces close to optimal results in CL, and even better results in terms of SC.

Our experiments over references and read datasets of large model organisms and bacterial pangenomes show that the CL decreases by up to 27% and the SC by up to 91% over UST^[6]^. Compared to unitigs, the CL decreases by up to 60% and SC by up to 97%. These improvements come often at just minor costs, as computing our small representation (which includes a run of BCALM2) takes less than twice as long than computing unitigs with BCALM2, and takes less than 35% longer in most cases. Even if the memory requirements for large read datasets increase, they stay within the limits of a modern server.

Finally we show that besides the smaller size of a minimum SPSS, it also has advantages in downstream applications. As an example of a kmer-based method, we query our compressed representation with the tools SSHash [45] and Bifrost [23]. These are state-of-the-art tools supporting kmer-based queries in genomic sequences, using a representation of a kmer set as a set of unitigs. By simply replacing unitigs with the strings produced by our greedy heuristic, and without modifications to Bifrost and a minor modification to SSHash disabling features that require unique kmers, we get a speedup of up to 4.26× over unitigs, and up to 2.10× over strings computed by UST and ProphAsm. We call the modified version of SSHash “SSHash-Lite”.

## 2 Results

### 2.1 Basic graph notation

We give detailed definitions for our notation below in Section 5.1, but give an intuition about the required notation for the results section here already. Note that our notation deviates from standard mathematical bidirected graph notations, but it is useful in practice as it allows to implement bidirected graphs on top of standard graph libraries. We assume that the reader is familiar with the general concept of de Bruijn graphs.

Our bidirected graphs are *arc-centric bidirected de Bruijn graphs*. Arc-centric de Bruijn graph means that kmers are on the arcs, and nodes represent *k* – 1 overlaps between kmers. We represent the bidirected graph as doubled graph, i.e. by having a separate forward and reverse arc for each kmer and a separate forward and reverse node for each *k* – 1 overlap. In this graph, *binodes* are ordered pairs (*v*, *v*^−1^) of nodes that are reverse complements of each other, and *biarcs* are ordered pairs (*e*, *e*^−1^) of arcs that are reverse complements of each other. Two biarcs (*e*, *e*^−1^) and (*f*, *f*^−1^) are consecutive if the normal arcs e and *f* are consecutive, i.e. the *f* leaves the node entered by *e*. A *biwalk* is a sequence of consecutive biarcs. If a biwalk visits a biarc, then it is considered to be covering both directions of the biarc. See Figure 3a for an example.

### 2.2 Matchtigs as a minimum plain text representation of kmer sets

We introduce the *matchtig algorithm* that computes a character-minimum SPSS for a set of genomic sequences. While former heuristics (ProphAsm, UST) did not allow to repeat kmers, our algorithm explicitly searches for all opportunities to reduce the character count in the SPSS by repeating kmers. Consider for example the arc-centric de Bruijn graph in Figure 1a. When representing its kmers without repetition as in Figure 1b, we need 43 characters and 7 strings. But if we allow to repeat kmers as in Figure 1d, we require only 39 characters and 5 strings. It turns out that structures similar to this example occur often enough in real genome graphs to yield significant improvements in both character and string count of an SPSS.

**Figure 1:**
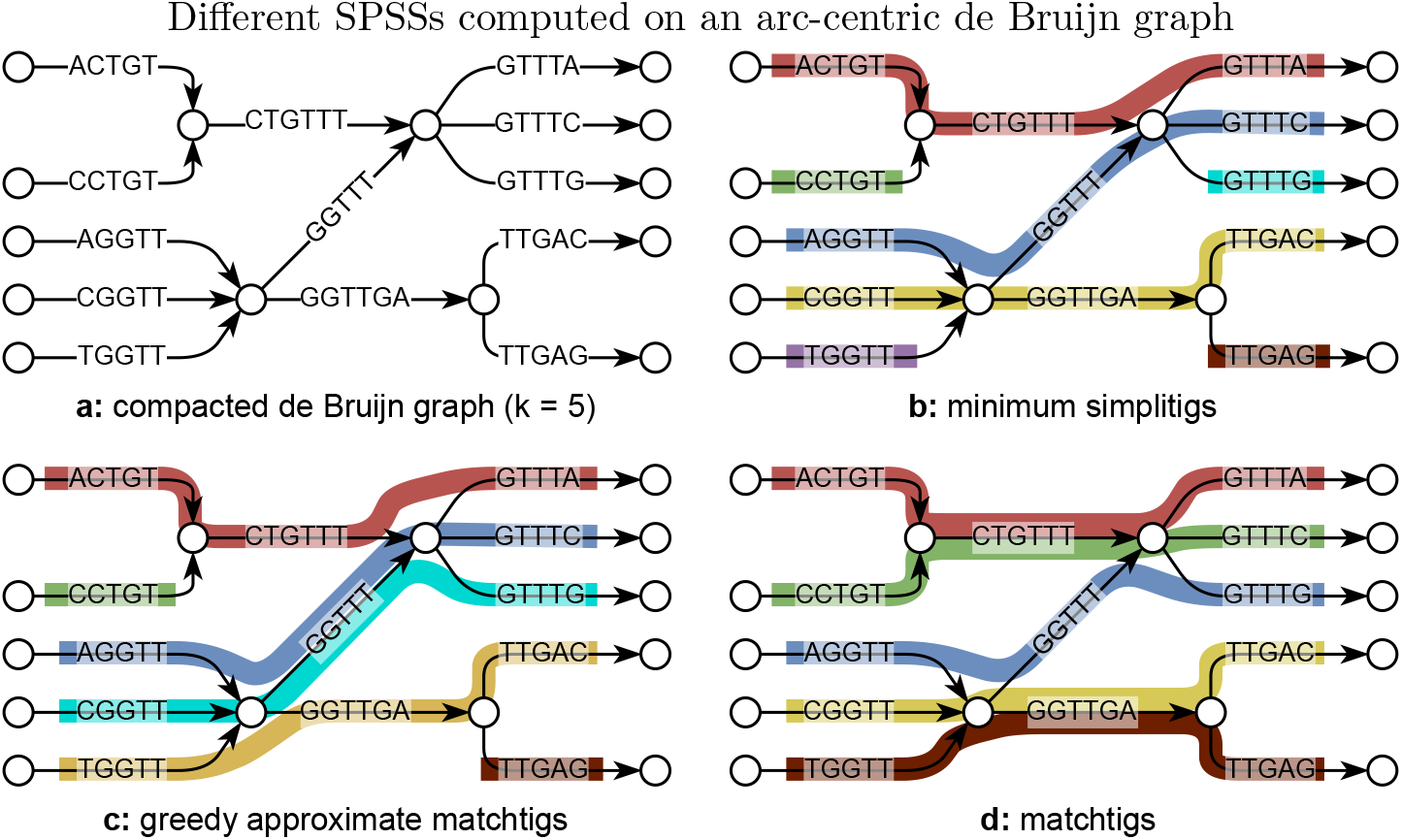
*k* = 5. For simplicity, the reverse complements of all nodes and arcs are omitted. **a:** the original de Bruijn graph in compacted form has 13 unitigs with 70 total characters. **b:** example of simplitigs with 7 strings and 43 total characters. **c:** example of greedily approximated matchtigs with 6 strings and 40 total characters. **d:** example of matchtigs with 5 strings and 39 total characters.

Similar to previous heuristics, our algorithm works on the compacted bidirected de Bruijn graph of the input sequences. However, we require an arc-centric de Bruijn graph, but this can be easily constructed from the node-centric variant (see Section 5.3). In this graph we find a min-cost circular biwalk that visits each biarc at least once, and that can jump between arbitrary nodes at a cost of *k* – 1. This formulation is very similar to the classic Chinese postman problem [53], formulated as follows: find a min-cost circular walk in a directed graph that visits each arc at least once. This similarity allows us to adapt a classic algorithm from Edmonds and Johnson that solves the Chinese postman problem [54] (the same principle was applied in [55]). They first reduce the problem to finding a min-cost Eulerisation via a min-cost flow formulation, and then further reduce that to min-cost perfect matching using a many-to-many min-cost path query between unbalanced nodes. In a similar work [56], the authors solve the Chinese postman problem in a bidirected de Bruijn graph by finding a min-cost Eulerisation via a min-cost flow formulation. As opposed to [54, 55] and us, in [56] the authors propose to solve the min-cost flow problem directly with a min-cost flow solver. We believe this to be infeasible for our problem, since the arbitrary jumps between nodes require the graph in the flow formulation to have a number of arcs quadratic in the number of nodes.

Our resulting algorithm is polynomial but while it runs fast for large bacterial pangenomes, it proved practically infeasible to build the matching instance for very large genomes (≥ 500Mbp). This is because each of the min-cost paths found translates into roughly one edge in the matching graph, and the number of min-cost paths raises quadratically if the graph gets denser. Thus, our algorithm ran out of memory when constructing it for larger genomes, and for those where we were able to construct the matching instance, the matcher itself suffered from integer overflows, since it uses 32-bit integers to store the instance. Hence, for practical purposes, we introduce a greedy heuristic to compute approximate matchtigs. This heuristic does not build the complete instance of the matching problem, but just greedily chooses the shortest path from each unbalanced node to Eulerise the graph. This reduces the amount of paths per node to at most one, and as a result, the heuristic uses significantly less memory, runs much faster, and achieves near optimal speedups when run with multiple threads (see Additional file 5). While it can in theory produce suboptimal results as in Figure 1c, in practice, the size of the greedily computed strings is very close to that of matchtigs, and the number of strings is always smaller.

Moreover, the minimality of matchtigs allows us to exactly compare, for the first time, how close heuristic algorithms to compute simplitigs are to optimal SPSS (on smaller genomes and on bacterial pangenomes, due to the resource-intensiveness of optimal matchtigs).

Our implementations are available^[7]^ as both a library and a command line tool, both written in Rust. They support both GFA and fasta file formats with special optimisations for fasta files produced by BCALM2 or GGCAT. Additionally, our implementations support gzip-compressed input and output, as well as outputting an ASCII-encoded bitvector of duplicate kmers.

### 2.3 Compression of model organisms

We evaluate the performance of our proposed algorithms on three model organisms: *C. elegans*, *B. mori* and *H. sapiens*. We benchmark the algorithms on both sets of short reads (average length 300 for C. elegans and B. mori, and 296 for H. sapiens) and reference genomes of these organisms. On human reads, we filter the data during processing so that we keep only kmers that occur at least 10 times (min abundance = 10).

We use the metrics cumulative length (CL) and string count (SC) as in [43]. The CL is the total number of characters in all strings in the SPSS, and the SC is the number of strings. We evaluate our algorithms against the same large genomes as in [43], using both the reference genome and a full set of short reads of the respective species (see Table 1 for the results). Since UST as well as matchtigs and greedy matchtigs require unitigs as input, and specifically UST needs some extra information in a format only output by BCALM2 [33], we run BCALM2 to compute unitigs from the input strings. We chose *k* = 31, as it is commonly used in kmer-based methods. While for larger genomes, larger *k* are used as well, we use the value *k* = 31 throughout the main matter to allow for easier comparison between results. Further, for all data sets but the C. elegans reference the matchtigs algorithm ran out of the given 256GiB memory, so we only compute greedy matchtigs for those.

**Table 1:** Quality and performance of compressing model organisms

| genome | algorithm | CL ratio | SC ratio | time [s] | memory [GiB] |
| --- | --- | --- | --- | --- | --- |
| C. elegans<br>(reads) | B2 | 1.00 | 1.00 | 2402 | 5.54 |
|  | B2+UST | 0.58 | 0.37 | 3424 (1.43) | 17.6 (3.18) |
|  | ProphAsm | 0.55 | 0.34 | 5433 | 56.5 |
|  | B2+GREEDY | 0.45 (0.79) | 0.11 (0.28) | 3057 (1.27) | 41.0 (7.41) |
| B. mori<br>(reads) | B2 | 1.00 | 1.00 | 6406 | 9.95 |
|  | B2+UST | 0.55 | 0.35 | 9896 (1.54) | 56.2 (5.64) |
|  | ProphAsm | 0.52 | 0.31 | 27912 | 157 |
|  | B2+GREEDY | 0.41 (0.74) | 0.06 (0.18) | 11793 (1.84) | 123 (12.4) |
| H. sapiens<br>(reads) | B2 | 1.00 | 1.00 | 168938 | 12.4 |
|  | B2+UST | 0.67 | 0.46 | 170427 (1.01) | 29.0 (2.34) |
|  | B2+GREEDY | 0.57 (0.84) | 0.22 (0.48) | 209646 (1.24) | 68.5 (5.52) |
| C. elegans | B2 | 1.00 | 1.00 | 52.7 | 0.96 |
|  | B2+UST | 0.92 | 0.34 | 58.6 (1.11) | 0.96 (1.00) |
|  | ProphAsm | 0.92 | 0.30 | 133 | 3.78 |
|  | B2+GREEDY | 0.90 (0.98) | 0.06 (0.18) | 59.9 (1.14) | 0.96 (1.00) |
|  | B2+MATCH | 0.90 (0.98) | 0.07 (0.23) | 380 (7.21) | 1.34 (1.40) |
| B. mori | B2 | 1.00 | 1.00 | 244 | 1.92 |
|  | B2+UST | 0.78 | 0.34 | 303 (1.24) | 1.92 (1.00) |
|  | ProphAsm | 0.76 | 0.28 | 716 | 13.8 |
|  | B2+GREEDY | 0.72 (0.92) | 0.06 (0.19) | 334 (1.37) | 2.42 (1.26) |
| H. sapiens | B2 | 1.00 | 1.00 | 1787 | 6.29 |
|  | B2+UST | 0.79 | 0.33 | 2249 (1.26) | 8.80 (1.40) |
|  | ProphAsm | 0.76 | 0.26 | 6677 | 130 |
|  | B2+GREEDY | 0.71 (0.91) | 0.03 (0.10) | 4999 (2.80) | 17.3 (2.75) |

We chose $k = 31$ and a min abundance of 10 for Homo sapiens reads and 1 for all others. The CL and SC ratios are between compressed strings and unitigs, and in parentheses are the ratios between our algorithm and the best competitor. B2 means BCALM2. For time and memory, we report the total time and maximum memory required to compute the tigs from the respective data set. BCALM2 directly computes unitigs and ProphAsm directly computes heuristic simpltigs. UST, GREEDY and MATCH compute heuristic simpltigs, greedy matchtigs and matchtigs from unitigs. The number in parentheses behind time and memory indicates the slowdown/increase over computing just unitigs with BCALM2. All algorithms were run with 28 threads, except for UST which supports only one thread (the preceding run of BCALM2 was still executed with 28 threads), and ProphAsm, which supports only one thread as well. Matchtigs are too expensive to run on all genomes except for the C. elegans reference, and ProphAsm takes too much time on H. sapiens reads, especially since it does not support minimum abundance. The lengths of the genomes are 100Mbp for C. elegans, 482Mbp for B. mori and 3.21Gbp for H. sapiens and the read data sets have a coverage of 64x for C. elegans, 58x for B. mori and 300x for H. sapiens.

On read data sets where we keep all kmers, our greedy heuristic achieves an improvement of up to 27% CL and 82% SC over the best competitor (UST-tigs). The human read data set has smaller improvements, however it was processed with a min abundance of 10, yielding longer unitigs with less potential for compression. On reference genomes the improvement in CL is smaller with up to 7%, however the improvement in SC is much larger with up to 91%.

For C. elegans, where computing matchtigs is feasible as well, we observe that they yield no significant improvement in CL, but are even slightly worse in SC than the greedy heuristic. The greedy heuristic actually optimises SC more than the optimal matchtigs algorithm. That is because the matching instance in the optimal algorithm is built to optimise CL, and whenever joining two strings does not alter CL, the choice is made arbitrarily. On the other hand, the greedy heuristic makes as many joins as possible, as long as a join does not worsen the CL. This way, the greedy heuristic actually prioritises joining two strings even if it does not alter the CL. For more details, see Sections 5.6 and 5.8. See Additional file 1 for more quality measurements with different kmer size and min. abundance.

We assume that the improvements correlate inversely with the average length of maximal unitigs of the data set. Our approach achieves a smaller representation by joining unitigs with overlapping ends, avoiding the repetition of those characters. This has a natural limit of saving at most *k* – 1 characters per pair of unitigs joint together, so at most *k* – 1 characters per unitig. In turn, the maximum fraction of characters saved is bound by *k* – 1 divided by the average length of unitigs. In Additional file 1 we have varied the kmer size and min. abundance for our data sets to vary the average length of unitigs. This gives us visual evidence for a correlation between average unitig length and decrease in CL.

Our improvements come at often negligible costs in terms of time and memory. Even for read sets, the run time at most doubles compared to BCALM2 in the worst case. However, the memory consumption rises significantly for read sets. This is due to the high number of unitigs in those graphs and the distance array of Dijkstra’s algorithm, whose size is linear in the number of nodes and the number of threads. See Additional file 2 for more performance measurements with different kmer size and min. abundance.

### 2.4 Compression of pangenomes

In addition to model organisms with large genomes, we evaluate our algorithms on bacterial pangenomes of *N. gonorrhoeae*, *S. pneumoniae*, *E. coli* and *Salmonella*, as well as a *human* pangenome. We use the same metrics as for model organisms. For the bacterial genomes, we choose *k* = 31, but also for the human genome for the reasons argued above, and also for easier comparability of the results on the different genomes. We show the results in Table 2. See Additional file 3 for more quality measurements with different kmer size and min. abundance, and Additional file 4 for more performance measurements with different kmer size and min. abundance. In neither of them we have included Salmonella or human, as they take too much time.

**Table 2:** Quality and performance of compressing pangenomes

| pangenome | algorithm | CL ratio | SC ratio | time [s] | memory [MiB] |
| --- | --- | --- | --- | --- | --- |
| 1102x<br>N. gonorrhoeae | B2 | 1.00 | 1.00 | 29.1 | 4351 |
|  | B2+UST | 0.63 | 0.35 | 31.1 (1.07) | 4351 (1.00) |
|  | ProphAsm | 0.62 | 0.33 | 735 (25.3) | 207 (0.05) |
|  | B2+GREEDY | 0.57 (0.93) | 0.18 (0.54) | 30.2 (1.04) | 4351 (1.00) |
|  | B2+MATCH | 0.57 (0.92) | 0.18 (0.56) | 31.1 (1.07) | 4351 (1.00) |
| 616x<br>S. pneumoniae | B2 | 1.00 | 1.00 | 26.1 | 3146 |
|  | B2+UST | 0.61 | 0.35 | 31.1 (1.19) | 3146 (1.00) |
|  | ProphAsm | 0.60 | 0.33 | 445 (17.0) | 434 (0.14) |
|  | B2+GREEDY | 0.53 (0.89) | 0.13 (0.41) | 29.0 (1.11) | 3146 (1.00) |
|  | B2+MATCH | 0.52 (0.88) | 0.14 (0.44) | 41.8 (1.60) | 3146 (1.00) |
| 3682x<br>E. coli | B2 | 1.00 | 1.00 | 334 | 7117 |
|  | B2+UST | 0.60 | 0.35 | 417 (1.25) | 7117 (1.00) |
|  | ProphAsm | 0.59 | 0.32 | 13339 (39.9) | 7221 (1.01) |
|  | B2+GREEDY | 0.51 (0.87) | 0.11 (0.33) | 384 (1.15) | 7117 (1.00) |
|  | B2+MATCH | 0.50 (0.85) | 0.12 (0.37) | 861 (2.58) | 7962 (1.12) |
| ~309kx<br>Salmonella | B2 | 1.00 | 1.00 | 82417 | 13007 |
|  | B2+UST | 0.57 | 0.36 | 82841 (1.01) | 13007 (1.00) |
|  | B2+GREEDY | 0.46 (0.81) | 0.11 (0.30) | 82726 (1.00) | 19512 (1.50) |
| 2505x<br>Human | CF | 1.00 | 1.00 | 77582 | 411472 |
|  | CF+ProphAsm | 0.68 | 0.31 | 82797 (1.07) | 411472 (1.00) |
|  | CF+GREEDY | 0.63 (0.93) | 0.16 (0.50) | 83507 (1.08) | 411472 (1.00) |

We chose $k = 31$ and a min abundance of 1. The CL and SC ratios are between compressed strings and unitigs, and in parentheses are the ratios between our algorithm and the best competitor. B2 means BCALM2. For time and memory, we report the total time and maximum memory required to compute the tigs from the respective data set. BCALM2 directly computes unitigs and ProphAsm directly computes heuristic simpltigs. UST, GREEDY and MATCH compute heuristic simpltigs greedy matchtigs and matchtigs from unitigs. The number in parentheses behind time and memory indicates the slowdown/increase over computing just unitigs with BCALM2. All algorithms were run with 28 threads, except for UST which supports only one thread (the preceding run of BCALM2 was still executed with 28 threads), and ProphAsm, which supports only one thread as well. The N. gonorrhoeae pangenome contains 8.36 million unique kmers, the S. pneumoniae pangenome contains 19.3 million unique kmers, the E. coli pangenome contains 341 million unique kmers, the Salmonella pangenome contains 657 million unique kmers and the human pangenome contains 2.8 billion unique kmers. Due to its size, ProphAsm and MATCH could not be run on the Salmonella pangenome. Also due to size, BCALM2 did not run on the human pangenome, hence we used Cuttlefish 2. To still be able to compare against competitors, we ran ProphAsm on the unitigs produced by Cuttlefish 2 (UST requires extra information from BCALM2).

Our algorithms improve CL up to 19% (using greedy matchtigs) over the best competitor and SC up to 70% (using greedy matchtigs). Matchtigs always achieve a slightly lower CL and slightly higher SC than greedy matchtigs, but the CL of greedy matchtigs is always at most 2% worse than that of matchtigs. We again assume that the improvements are correlated inversely to the average size of unitigs, as suggested by the experiments in Additional file 3. These improvements come at negligible costs, using at most 23% more time and 11% more memory than BCALM2 when computing greedy matchtigs, except for the large salmonella pangenome, which took 50% more memory. The higher memory consumption is due to the graph being more tangled due to the high number of genomes in the pangenome. For matchtigs, the time increases by less than a factor of two and memory by at most 12% compared to BCALM2.

### 2.5 Kmer-based short read queries

Matchtigs have further applications beyond merely reducing the size required to store a set of kmers. Due to their smaller size and lower string count, they can make downstream applications more efficient. To make a concrete example, in this section we focus on *membership queries*. As already explained, each SPSS (unitigs, UST-tigs, matchtigs, etc.) can be considered as a (multi-) set of kmers. Given a kmer, a membership query is to verify whether the kmer belongs to the set or not. We focus on *exact* queries, rather than approximate, i.e., if a kmer does not belong to the set then the answer to the query *must* be “false”. Assessing the membership to the set for a string *Q* longer than *k* symbols is based on the answers to its constituent kmers: only if *at least* ⌊*θ* × (|*Q*| – *k* + 1)⌋ kmers of *Q* belongs to the set, then *Q* is considered to be present in the set. The threshold *θ* is therefore an “inclusion” rate, which we fix to 0.8 for the experiments in this section.

To support fast membership queries in compressed space, we build an SSHash-Lite^[8]^dictionary over each SPSS. SSHash-Lite is a relaxation of SSHash [45, 57] in that it supports membership queries *without* requiring each kmer to appear once in the underlying SPSS. In short, SSHash is a compressed dictionary for kmers – based on minimal perfect hashing [58] and minimizers [59] – which, for an input SPSS without duplicates and having *n* (distinct) kmers, assigns to each kmer in the input a unique integer number from 0 to *n* – 1 by means of a Lookup query. The result of Lookup for any kmer that is *not* contained in the input SPSS is –1. Therefore, SSHash serves the same purpose of a minimal perfect hash function over a SPSS but it is also able to reject alien kmers. Two variants of SSHash were proposed – a *regular* and a *canonical* one. The canonical variant uses some extra space compared to the regular one but queries are faster to answer. (For all further details, we point the reader to the original papers [45, 57].)

Now, to let SSHash be able to query SPSSs with possible duplicate kmers (e.g., matchtigs), it was only necessary to modify the return value of the Lookup query to just return “true” if a kmer if found in the dictionary rather than its unique integer identifier (respectively, “false” if a kmer is not found instead of –1). Therefore, SSHash-Lite can be directly used to index and query the unitigs, UST-tigs, and matchtigs as well.

We compare the performance of SSHash-Lite when indexing unitigs, UST-tigs, and matchtigs in Table 3. We build the SPSSs from three datasets: a ~309kx Salmonella Enterica pangenome; a 300 × coverage human short read dataset filtered to exclude kmers with an abundance lower than 10; and a 2505x human pangenome. The Salmonalla pangenome was queried with 3 million random Salmonella short reads with lengths between 70 and 502, and an N75 of 302. The human queries for both the human read dataset and the human pangenome are 3 million random short reads (296 bases each) from the human read dataset.

**Table 3:** Performance characteristics of querying different tigs with SSHash-Lite.

| (a) regular SSHash-Lite |  |  |  |  |  |  |
| --- | --- | --- | --- | --- | --- | --- |
| genome | algorithm | index time<br>[min] | search time<br>[sec] | search<br>speedup | index size<br>[GiB] | size imprv. |
| ~309kx<br>Salmonella<br>(0.75) | unitigs | 2.77 | 3027 | 1.00 | 1.04 | 1.00 |
|  | UST | 2.42 | 1491 | 2.03 | 0.71 | 1.48 |
|  | gMatchtigs | 2.60 | 710 | 4.26 (2.10) | 0.65 | 1.60 (1.08) |
| Human<br>reads<br>(0.75) | unitigs | 18.9 | 558 | 1.00 | 4.60 | 1.00 |
|  | UST | 17.1 | 499 | 1.12 | 3.63 | 1.27 |
|  | gMatchtigs | 19.2 | 384 | 1.45 (1.30) | 3.47 | 1.33 (1.05) |
| 2505x<br>Human<br>(0.65) | unitigs | 15.1 | 515 | 1.00 | 3.63 | 1.00 |
|  | ProphAsm | 14.0 | 421 | 1.22 | 2.86 | 1.27 |
|  | gMatchtigs | 14.8 | 363 | 1.42 (1.16) | 2.86 | 1.27 (1.00) |

| (b) canonical SSHash-Lite |  |  |  |  |  |  |
| --- | --- | --- | --- | --- | --- | --- |
| genome | algorithm | index time<br>[min] | search time<br>[sec] | search speedup | index size<br>[GiB] | size imprv. |
| ~309kx<br>Salmonella<br>(0.75) | unitigs | 3.94 | 1576 | 1.00 | 1.13 | 1.00 |
|  | UST | 3.30 | 961 | 1.64 | 0.78 | 1.44 |
|  | gMatchtigs | 3.71 | 572 | 2.75 (1.68) | 0.74 | 1.52 (1.06) |
| Human<br>reads<br>(0.75) | unitigs | 25.0 | 373 | 1.00 | 5.02 | 1.00 |
|  | UST | 23.4 | 324 | 1.15 | 4.05 | 1.24 |
|  | gMatchtigs | 26.3 | 266 | 1.40 (1.22) | 3.94 | 1.28 (1.03) |
| 2505x<br>Human<br>(0.65) | unitigs | 21.3 | 340 | 1.00 | 4.26 | 1.00 |
|  | ProphAsm | 20.1 | 258 | 1.32 | 3.48 | 1.22 |
|  | gMatchtigs | 21.1 | 232 | 1.46 (1.11) | 3.52 | 1.21 (0.99) |

SSHash-Lite is run with $k = 31$ and a kmer-inclusion rate of 0.8. On the Salmonella pan-genome we used a minimizer length of 17 for the regular index and a minimizer length of 16 for the canonical index. On the human reads we used a minimizer length of 20 for the regular index and a minimizer length of 19 for the canonical index. On the human pangenome we used a minimizer length of 19 for the regular index and a minimizer length of 20 for the canonical index. The search speedup is with respect to unitigs, and the search speedup in parentheses is with respect to the strings computed by UST. Index time is the end-to-end time required to build the SSHash-Lite index: it includes reading the collections from disk and building the data structure using external memory. Searching time is the time required to check which reads have at least 80% of their kmers in the input SPSS. The number in parentheses under the genome is the kmer hitrate, i.e. the fraction of kmers from the query that are part of the queried dataset.

We see that matchtigs improve the performance of membership queries in *both space and time* compared to unitigs and UST-tigs. While the difference is more evident when compared to unitigs, matchtigs also consistently outperform UST-tigs - achieving the lowest space usage and faster query time across almost all combinations of dataset and index variant (regular/canonical).

Note again that the speed up in searching time is more evident on the human reads dataset since it is much larger than the Salmonella pan-genome and it is generally less evident for the canonical index variant of SSHash-Lite because it is approximately 2× faster to query than the regular one. Remarkably, regular SSHash-Lite over matchtigs achieves 27 – 60% reduction in space over unitigs while being also 4.26× faster to query on the human reads datasets. Compared to UST-tigs instead, matchtigs still retain 2.10× faster query time while improving space by up to 8%. These results were achieved on a typical bioinformatics compute node with many logical cores (256) and a large amount of RAM (2TB). In Additional file 10 we performed the same experiment on a server with focus on single-thread performance, achieving slightly smaller improvements.

The reduction in index space when indexing matchtigs is to be attributed to the lower string count and fewer nucleotides in the collection. The speedups achieved by SSHash-Lite when indexing matchtigs instead of unitigs can be explained as follows. When querying, SSHash-Lite streams through the kmers of the query. At the beginning, the tig containing the first kmer of the query is determined using a minimal perfect hash function over the minimizers of the input SPSS, as well as the position of the kmer in the tig. For the subsequent kmers of the query, SSHash-Lite attempts to “extend” the matching of the kmer against the identified tig by just comparing the single nucleotide following the previous kmer in the tig. Extending a match in this way is extremely fast not only because just a single nucleotide needs to be compared but also because it is a very cache-friendly algorithm, dispensing random accesses to the index entirely. However, each time an extension is not possible (either because we have a mismatch or we have reached the end of the current tig) a “full” new search is made in the index. The search consists in evaluating the minimal perfect hash function and locating the kmer inside another tig. Clearly, a search is much more expensive due to cache misses compared to an extension. Now, using longer tigs with a lower tig count - the case for the matchtigs – increases the chance of extension, or equivalently, decreases the number of full searches in the index. Compared to UST-tig, matchtigs can be faster to query exactly because allowing repeated kmers to appear in the tigs further helps in creating opportunities for extension. Therefore, by reducing the number of full searches, we can reduce the overall runtime of the query.

## 3 Discussion

Kmer-based methods have found wide-spread use in many areas of bioinformatics over the past years. However, they usually rely on unitigs to represent the kmer sets, since they can be computed efficiently with standard tools [33, 23, 40, 41]. Unitigs have the additional property that the de Bruijn graph topology can easily be reconstructed from them, since they do not contain branching nodes other than on their first and last kmer. However, this property is not usually required by kmer-based methods, which has opened the question if a smaller set of strings other than unitigs can be used to represent the kmer sets. If such a representation was in plain text, it should be usable in most kmer-based tools, by simply feeding it to the tool instead of unitigs.

Previous work has relaxed the unitig requirement of the representation of the kmer sets to arbitrary strings without kmer repetitions. This resulted in a smaller representation, leading to improvements in downstream applications. Additionally, previous work considered whether that finding an optimal representation without repeated kmers is NP-hard, which was then disproven and shown to be linear-time solvable. We have shown that by allowing repetitions, there is a polynomial optimal algorithm that achieves better compression and improvements in downstream applications.

## 4 Conclusions

Our *optimum* algorithm compresses the representation significantly more than previous work. For practical purposes we also propose a greedy heuristic that achieves near-optimum results, while being suitable for practical purposes in runtime and memory. Specifically, our algorithms achieve a decrease of 27% in size and 91% in string count over UST. Additionally we have shown that our greedy representation speeds up downstream applications, giving an example with a factor of 2.10 compared to previous compressed representations.

Our implementation is available as a stand-alone command-line tool and as a library. We hope that our efficient algorithms result in a wide-spread adoption of near-minimum plain-text representations of kmer sets in kmer-based methods, resulting in more efficient bioinformatics tools.

## 5 Methods

We first give some preliminary definitions in Section 5.1 and define our problem in Section 5.2. Note that to stay closer to our implementation, our definitions of bidirected de Bruijn graphs differ from those in e.g. [56]. However, the concepts are fundamentally the same. Then in Sections 5.3 to 5.7 we describe how to compute matchtigs. The whole algorithm is summarised by an example in Figure 2. For simplicity, we describe the algorithm using an uncompacted de Bruijn graph. However, in practice it is much more efficient to use a compacted de Bruijn graph, but our algorithm can be adapted easily: simply replace the costs of 1 for each original arc with the number of uncompacted arcs it represents. In Section 5.8 we describe the greedy heuristic.

**Figure 2:**
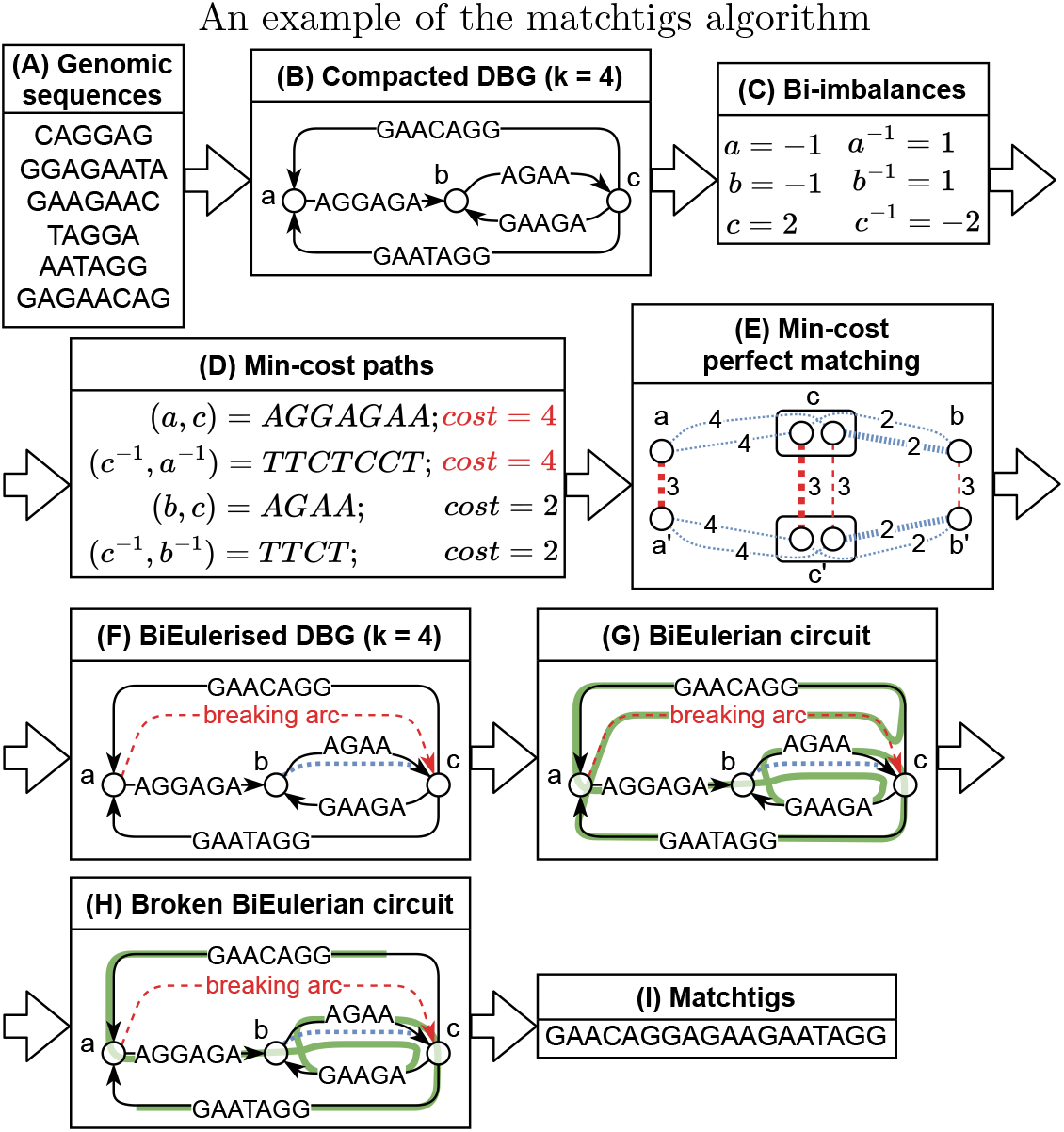
**(A)** The input genomic sequences. **(B)** We first build an arc-centric compacted de Bruijn graph (for simplicity, the reverse complements of the nodes and arcs are not shown) **(C)** In the graph we compute the bi-imbalances of the nodes (the difference between outdegree and indegree). **(D)** From each node with negative bi-imbalance we compute the min-cost paths to all reachable nodes with positive bi-imbalance. The costs of each arc are the amount of characters required to join two strings from the negative to the positive node while repeating the kmers between the nodes. Specifically, the costs of an arc are |*s*| – (*k* – 1), where |*s*| is the length of its label. **(E)** Using a min-cost perfect matching instance built from the min-cost paths, we decide which bi-imbalances should be fixed by repeating kmers. The blue/tightly dashed edges are joining edges stemming from the min-cost paths. The red edges in longer dashes indicate that a node should stay unmatched, i.e. that fixing its bi-imbalance requires breaking arcs. The solution edges are highlighted in bold. There is one node in the matching problem for each binode in the original graph. The nodes *x’* are not reverse complements of nodes *x*, but stem from a reduction that makes a copy of each node. For more details, refer to Section 5.6. **(F)** For each joining edge in the solution we insert a joining arc into the DBG (in blue, small dashes), always directed such that the overall bi-imbalance decreases. The remaining imbalance is removed by inserting arbitrary breaking arcs (in read, longer dashes). **(G)** We compute a biEulerian circuit in the balanced graph. **(H)** We break the biEulerian circuit at all breaking arcs. **(I)** We output the strings spelled by the broken walks.

**Figure 3:**
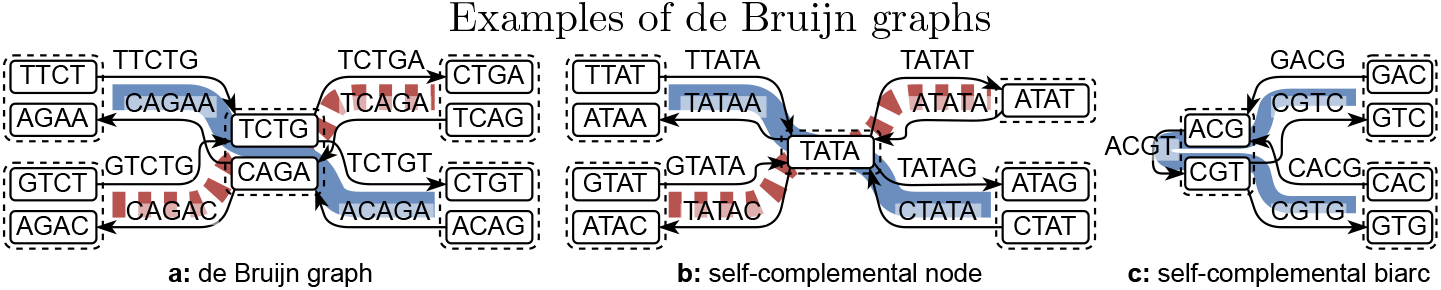
Binodes are surrounded by a dashed box, where self-complemental binodes contain only one graph node. **a:** a de Bruijn graph of the strings TTCTGA and GTCTGT. The colored/patterned lines are an arc-covering set of biwalks. **b:** a de Bruijn graph containing two self-complemental nodes. The colored/patterned lines are a set of biwalks that visit each biarc exactly once. **c:** a de Bruijn graph containing a self-complemental biarc. The colored line is a biwalk that visits each biarc exactly once.

### 5.1 Preliminaries

We are given an alphabet Γ and all strings in this work have only characters in Γ. Further, we are given a bijection comp: Γ → Γ. The *reverse complement* of a string *s* is s^−1^:= rev(comp*(*S*)) where rev denotes the reversal of a string and comp* the character-wise application of comp. For an integer *k*, a string of length *k* is called a *kmer*. From here on, we only consider strings of lengths at least *k*, i.e. strings that have at least one kmer as substring. We denote the prefix of length *k* – 1 of a kmer s by pre(s) and its suffix of length *k* – 1 by suf(*s*). The *spectrum* of a set of strings *S* is defined as the set of all kmers and their reverse complements that occur in at least one string *s* ∈ *S*, formally spec_*k*_(*S*):= {*r* ∈ Γ^*k*^| ∃*s* ∈ *S*: *r* or *r*^−1^ is substring of *s*}.

An *arc-centric de-Bruijn graph* (or short *de-Bruijn graph*) DBG_k_(*S*) = (*V*, *E*) of order *k* of a set of strings *S* is defined as a standard directed graph with nodes *V*:= {*s* | *s* ∈ spec_*k*–1_(*S*)} and arcs *E*:= {(pre(*s*),suf(*s*)) | *s* ∈ spec_*k*_(*S*)}. On top of this, we use the following notions of bidirectedness. An ordered pair of reverse-complementary nodes [*v*, *v*^−1^] ∈ *V* × *V* is called a *binode* and an ordered pair of reverse-complementary arcs [(*a*, *b*), (*b*^−1^, *a*^−1^)] ∈ *E* × *E* is called a *biarc*. Even though these pairs are ordered, reversing the order still represents the same binode/biarc, just in the other direction. *A* node *v* is called *canonical* if *v* is lexicographically smaller than *v*^−1^, and an arc (*a*, *b*) is called *canonical* if the kmer corresponding to (*a*, *b*) is lexicographically smaller or equal to the kmer corresponding to (*b*^−1^, *a*^−1^). If an arc or a node is its own reverse-complement (called *self-complemental*), then it is written as biarc [(*a*, *b*)] or binode [*v*]. See Figure 3 for examples of different bigraphs.

Since de Bruijn graphs are defined as standard directed graphs, we use the following standard definitions. The set of incoming (outgoing) arcs of a node is denoted by *E*^−^(*v*) (*E*^+^(*v*)), and the indegree (outdegree) is *d*^−^(*v*):=|*E*^−^(*v*)| (*d*^+^(*v*):= |*E*^+^(*v*)|). A *walk* in a de Bruijn graph is a sequence of adjacent arcs (followed in the forward direction) and a *unitig* is a walk in which all inner nodes (nodes with at least two incident walk-arcs) have exactly one incoming and one outgoing arc. The length |*w*| of a walk *w* is the length of the sequence of its arcs (counting repeated arcs as often as they are repeated). A *compacted de-Bruijn graph* is a de Bruijn graph in which all maximal unitigs have been replaced by a single arc. A *circular walk* is a walk that starts and ends in the same node, and a *Eulerian circuit* is a circular walk that contains each arc exactly once. A graph that admits a Eulerian circuit is *Eulerian*.

Assuming the complemental pairing of nodes and arcs defined above, we can define the following bidirected notions of walks and standard de Bruijn graph concepts. *Biwalks* and *circular biwalks* are defined equivalently to walks, except that they are sequences of biarcs. A biwalk *w* in a de Bruijn graph spells a string spell(*w*) of overlapping visited kmers. That is, spell(*w*) is constructed by concatenating the string *a* from *w*’*s* first biarc [(*a*, *b*), (*b*^−1^,*a*^−1^)] (or [(*a, b*)]) with the last character of *b* of the first and all following biarcs. See Figure 3 for examples of bidirected de Bruijn graphs and notable special cases.

A bidirected graph is *connected*, if between each pair of distinct binodes [*u, u*^−1^], [*v,v*^−1^] that are not reverse complements of each other, there is a biwalk from [*u,u*^−1^] to [*v,v*^−1^], or from [*u,u*^−1^] to [*v*^−1^,*v*]. We assume that our graph is connected, as on multiple disconnected components, our algorithm can be executed on each component, yielding a minimum result.

### 5.2 Problem overview

We are given a set of input strings *I* where each string has length at least *k*, and we want to compute a minimum spectrum preserving string set, defined as follows.

#### Definition 1

(SPSS) *A* spectrum preserving string set (*or* SPSS) *of I is a set* S *of strings of length at least k such that* spec_*k*_ (*I*) = spec_k_(*S*), *i.e. both sets of strings contain the same kmers, either directly or as reverse complement*.

Note that our definition allows kmers and their reverse complements to be repeated in the SPSS, both in the same string and in different strings. This is the only difference to the definition of Rahman and Medvedev [44] and simplitigs [43], which can be formalised as a set of strings *S_simputigs_* of length at least k such that for each kmer in the spectrum, either itself or its reverse complement (but *not both*) occurs *exactly once* in *exactly one* string in *S_simplitigs_*.

#### Definition 2

(Minimum SPSS) *The* size ||*S*|| *of an SPSS S is defined as*

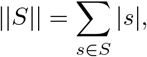

*where* |*s*| *denotes the length of string s. A* minimum *SPSS is an SPSS of minimum size*.

On a high level, our algorithm works as follows (see also Figure 2).

1. Create a bidirected de Bruijn graph from the input strings (see Section 5.3).
2. Compute the *bi-imbalances* of each node (see Section 5.5).
3. Compute the min-cost bipaths of length at most *k* – 1 from all nodes with negative bi-imbalance to all nodes with positive bi-imbalance (see Section 5.7).
4. Solve a min-cost matching instance with costs for unmatched nodes to choose a set of shortest bipaths with minimum cost by reduction to min-cost perfect matching (see Section 5.6).
5. *BiEulerise* the graph with the set of bipaths as well as arbitrary arcs between unmatched nodes.
6. Compute a *biEulerian circuit* in the resulting graph (see Section 5.5).
7. Break the circuit into a set of biwalks and translate them into a set of strings, which is the output minimum SPSS (see Section 5.4).

Note that in our implementation, a substantial difference is that we do not build the de Bruijn graph ourselves, but we expect the input to be a de Bruijn graph already. For our experiments, we use a compacted de Bruijn graph computed with BCALM2. We motivate the reasons for optimality while explaining our algorithm, but also give a formal proof in Additional file 8.

### 5.3 Building a compacted bidirected arc-centric de Bruijn graph from a set of strings

When building the graph we first compute unitigs from the input strings using BCALM2. Then we initialise an empty graph and do the following for each unitig:

1. We insert the unitig’s first *k* – 1-mer and its reverse complement as binode by inserting the two nodes separately and marking them as a bidirected pair, if it does not already exist. The existence is tracked with a hashmap, storing the two nodes corresponding to a kmer and its reverse complement if it exists.
2. We do the same for the last *k* – 1-mer of the unitig.
3. We add a biarc between the two binodes by inserting one forward arc between the forward nodes of the binodes, and one reverse arc between the reverse complement nodes of the binodes. The forward arc is labelled with the unitig, and the reverse arc is labelled with its reverse complement.

To save memory, we store the unitigs in a single large array, where each character is encoded as two-bit number. The keys of the hashmap and the labels of the arcs are pointers into the array, together with a flag for the reverse complement. Nodes do not need a label, as their label can be inferred from any of its incident arcs’ label. Recall that in the description of our algorithm, we use an uncompacted graph only for simplicity.

### 5.4 Reduction to the bidirected partial-coverage Chinese postman problem

We first compute the arc-centric de-Bruijn graph DBG_k_(*I*) of the given input string set *I* as described in Section 5.3. In DBG_k_(*I*), an SPSS S is represented by a *biarc-covering* set of biwalks *W* (the reverse direction of a biarc does not need to be covered separately). That is a set of biwalks such that each biarc is element of at least one biwalk (see Figure 3a). According to the definition of spell, the size of *S* is related to W as follows:

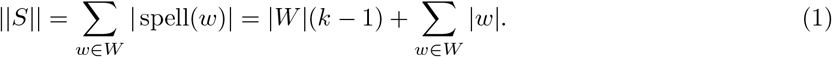

Each walk costs *k* – 1 characters because it contains the node *a* from its first biarc [(*a, b*), (*b*^−1^, *a*^−1^)] (or [(*a, b*)]), and it additionally costs one character per arc.

To minimise ||*S*||, we transform the graph as follows:

#### Definition 3

(Graph transformation) *Given an arc-centric de-Bruijn graph* DBG_*k*_(*I*) = (*V, E*), *the* transformed graph *is defined as* 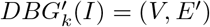k *where E’ is a multiset defined as E’*:= *E* ∪ (*V* × *V*). *In E’, arcs from E are marked as* non-breaking, *and arcs from V* × *V are marked as* breaking *arcs. The* cost function *c*(*e*), *e* ∈ *E*’ *assigns all non-breaking arcs the costs* 1 *and all breaking arcs the costs k* – 1.

The reverse complements of breaking arcs are breaking as well, and the same holds for non-breaking arcs. This means that biarcs always are either a pair of reverse complemental breaking arcs, in which case we call them *breaking biarcs*, or a pair of reverse complemental non-breaking arcs, in which case we call them *non-breaking biarcs*. By the same pattern, self-complemental biarcs are defined to be *breaking biarcs* or *non-breaking biarcs* depending on their underlying arc. Breaking arcs have the costs *k* – 1 because each breaking arc corresponds to starting a new walk, which by Equation (1) costs *k* – 1.

In the transformed graph we find a circular biwalk *w*\* of minimum cost that covers at least all original biarcs (to cover a biarc it is enough to traverse it once in one of its directions), as well as at least one breaking biarc. The reason for having at least one breaking biarc is that later we break the circular original-biarc-covering biwalk at all breaking biarcs to get a set of strings. If there was no breaking biarc, then we would get a circular string. Simply breaking a circular string at an arbitrary binode or a repeated non-breaking biarc does not produce a minimum solution in general^[9]^, hence we make it a part of the optimisation problem to find an optimal breaking point. We define such a walk as:

#### Definition 4

(Circular original-biarc-covering biwalk) *Given a transformed graph* 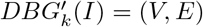k, *a* circular original-biarc-covering biwalk *is a circular biwalk* w *such that for each non-breaking arc (a, b) ∈ E there is a biarc* [(*a, b*), (*b*^−1^, *a*^−1^)], [(*b*^−1^, *a*^−1^), (*a, b*)] *or* [(*a, b*)] *in w. Additionally, w needs to contain at least one breaking biarc*.

#### Definition 5

(Costs of a circular original-biarc-covering biwalk) *Given a trans-formed graph* 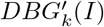k *and a circular original-biarc-covering walk w possibly consisting of biarcs* [(*a, b*), (*b*^−1^, *a*^−1^)] *and self-complemental biarcs* [(*a, b*)]. *The* costs *c*(*w*) *of w are the sum of the costs of each biarc and self-complemental biarc, where the costs of a biarc* [(*a, b*), (*b*^−1^, *a*^−1^)] *are c((a,b)), and the costs of a self-complemental biarc* [(*a, b*)] *are c*((*a,b*)).

This is similar to the directed Chinese postman problem (DCPP). In the DCPP, the task is to find a circular min-cost arc-covering walk in a directed graph. It is a classical problem, known to be solvable in *O*(*n*^3^) time [60] with a flow-based algorithm using e.g. [61] to compute min-cost flows. The partial coverage variant of the DCPP (requiring to cover only a subset of the arcs) is also known as the rural postman problem [62]. Further, the bidirected variant of the DCPP was discussed before in [56], and the authors also solved it using min-cost flow in *O*(*n*^2^log^2^(*n*)) time.

We break the resulting min-cost circular original-biarc-covering biwalk *w*\* at all breaking arcs (hence we require it to contain at least one breaking biarc, otherwise we would get a circular string). The returned SPSS is minimum, because the metric optimised when finding *w*\* matches Equation (1).

### 5.5 Solving the bidirected partial-coverage Chinese postman problem with min-cost integer flows

Edmonds and Johnson [54] introduced a polynomial-time flow-based approach that is adaptable to solve our variant of the DCPP. They show that finding a minimum circular arc-covering walk in a directed graph is equivalent to finding a minimum *Eulerisation* of the graph, and then any Eulerian circuit in the Eulerised graph. A Eulerisation is a multiset of arc-copies from the graph that makes the graph Eulerian if added^[10]^, and a minimum Eulerisation is one whose sum of arc costs is minimum among all such multisets. To find such a minimum cost set of arcs, they formulate a min-cost flow problem as an integer linear program as follows:

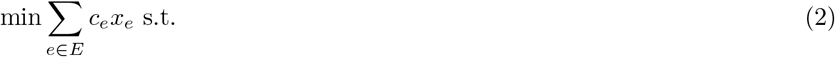

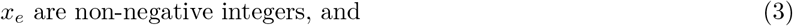

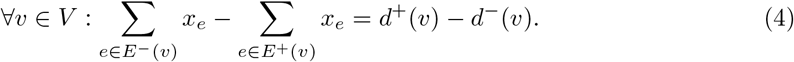

The variable *x_e_* is interpreted as the amount of flow through arc *e*, and the variable *c_e_* denotes the cost for assigning flow to an arc *e*. The costs are equivalent to the arc costs in the weighted graph specified by the DCPP instance. Objective (2) minimises the costs of inserted arcs as required. To ensure that the resulting flow can be directly translated into added arcs, Condition (3) ensures that the resulting flow is non-negative and integral. Lastly, Equation (4) is the *balance constraint*, ensuring that the resulting flow is a valid Eulerisation of the graph. This constraint makes nodes with missing outgoing arcs into *sources*, and nodes with missing incoming arcs into *sinks*, with demands matching the number of missing arcs. Note that in contrast to classic flow problems, this formulation contains no capacity constraint. For a solution of this linear program, the corresponding Eulerisation contains *x_e_* copies of each arc *e*.

To adapt this formulation to our variant of the DCPP, we need to make modifications, namely:

- compute the costs while treating self-complemental biarcs the same as other biarcs,
- allow for partial coverage,
- force cover at least one breaking arc (one of the arcs that are not required to be covered),
- adjust the balance constraint for biwalks and
- ensure that the resulting flow is *bidirected*, i.e. the flow of each arc equals the flow of its reverse complement.

#### Bidirected costs

If we would simply count the costs of each arc separately, then self-complemental biarcs would cost 1 for each repetition, while other biarcs would cost 2 for each repetition, since other biarcs consist for two arcs. To circumvent this, we only count arcs corresponding to canonical kmers in the cost function:

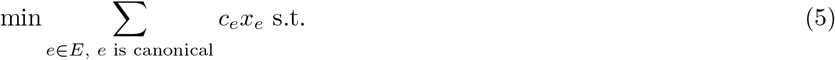

#### Partial coverage

In the partial coverage Chinese postman problem, we are additionally given a set *F* ⊆ *E* of arcs to be covered. In contrast to the standard DCPP, a solution walk only needs to cover all the arcs in *F*. In our case, the set *F* is the set of original arcs of the graph before Eulerisation. To solve the partial coverage Chinese postman problem we define outgoing covered arcs *F*^+^(*v*):= *F* ∩ E^+^(*v*), and incoming covered arcs *F*^−^(*v*):= *F* ∩ *E*^−^(*v*) for a node *v*, as well as the covered outdegree 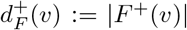k and the covered indegree 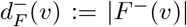k. Then we reformulate the balance constraint as:

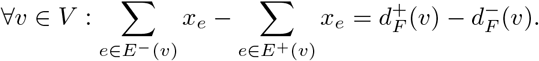

The resulting set of arcs is a minimum Eulerisation of the graph (*V, F*), and a Eulerian walk in this graph is equivalent to a minimum circular *F*-covering walk in the original graph.

#### Cover one breaking arc

Next to the partial coverage condition, we additionally require to cover at least one of the arcs that is not required to be covered. Since we forbid negative flows, we can express this as:

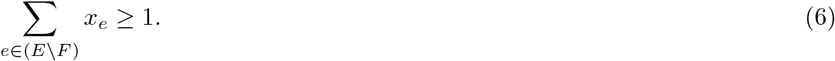

#### Bidirected balance

In contrast to Edmonds and Johnson, we are interested in a minimum circular *bi*walk that covers each original *bi*arc. Analogue to the formulation for directed graphs, we make the following definitions:

##### Definition 6

(BiEulerian circuits and graphs) *A* biEulerian circuit *in a bidirected graph is a bidirected circuit that visits each biarc exactly once. A* biEulerian graph *is a graph that admits a biEulerian circuit. A* biEulerisation *is a multiset of biarc-copies from the graph that makes a graph biEulerian if added. A* minimum biEulerisation *is one whose sum of arc costs is minimum among all biEulerisations of the same graph*.

We can compute a biEulerisation in the same way as we compute a Eulerisation, the only change is in the balance constraint. Observe that for a Eulerian graph, the *imbalance i_v_*:= *d*^−^(*v*) – *d*^+^(*v*) is zero for each node [63], because each node is entered exactly as often as it is exited. For binodes, the definition of the *bi-imbalance*bi_v_ of a binode [*v, v*^−1^] or [*v*] follows the same idea. However, in contrast to directed graphs, there are two (mutual exclusive^[11]^) special cases.

Binodes [*v, v*^−1^] ∈ *V* × *V* with *v* ≠ *v*^−1^ may have incident self-complemental arcs [(*v, v*^−1^)] and/or [(*v*^−1^, *v*)] (see Figure 3c for an example). If e.g. only [(*v, v*^−1^)] exists, then to traverse it, a biwalk needs to enter *v* twice. First, it needs to reach [*v, v*^−1^] via some biarc, and after traversing [(*v, v*^−1^)], it needs to leave [*v*^−1^, *v*] via a different biarc, whose reverse complement enters [*v, v*^−1^]. Hence, a self-complemental biarc alters the bi-imbalance of a node by two. See Figure 4 for an example of this situation. If only [(*v*^−1^, *v*)] exists, then the situation is symmetric. Therefore, for balance of [*v, v*^−1^], the self-complemental biarc [(*v, v*^−1^)] requires two biarcs entering [*v, v*^−1^] and the self-complemental biarc [(*v*^−1^, *v*)] requires two biarcs leaving *[v, v*^−1^]. If both self-complemental arcs exist (e.g. both [(*ATA, TAT*)] and [(*TAT, ATA*)] for a binode [*ATA, TAT*]), then a biwalk can traverse them consecutively from e.g. [*v, v*^−1^] by traversing first [(*v, v*^−1^)] and then [(*v*^−1^, *v*)], ending up in [*v, v*^−1^] again, such that the self-complemental arcs have a neutral contribution to the bi-imbalance. Resulting, the bi-imbalance of [*v, v*^−1^] is

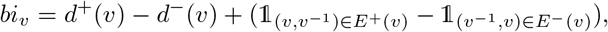

where 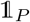k is 1 if the predicate *P* is true and 0 otherwise.

For self-complemental binodes [*v*] ∈ *V*, there is no concept of incoming or outgoing biarcs, since any biarc can be used to either enter or leave [*v*] (see Figure 3b for an example). Therefore, for balance, biarcs need to be paired arbitrarily, for which to be possible there needs to be an even amount of biarcs. If there is an odd amount, then there is one unpaired biarc, hence the bi-imbalance is 1. The following condition exerts this behaviour (using mod as the remainder operator):

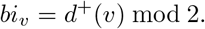

Finally, we include partial coverage to above bi-imbalance formulations by limiting the incoming and outgoing arcs to *F*. Further, to distinguish between self-complemental nodes and others, we denote the set of self-complemental nodes as *S* ⊆ *V* and the set of binodes that are not self-complemental as *T*:= *V* \ *S*. If self-complemental biarcs are included in the flow, then these alter the bi-imbalance by two, in the same way as they do in the equation of the bi-imbalance. We encode this on the left side of the equation. Then we get the following modified coverage constraint:

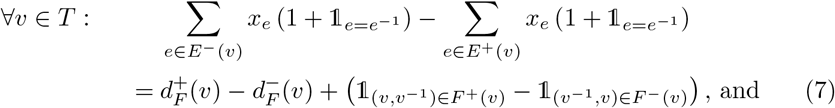

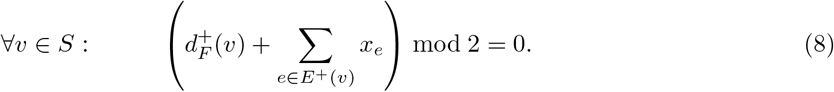

**Figure 4:**
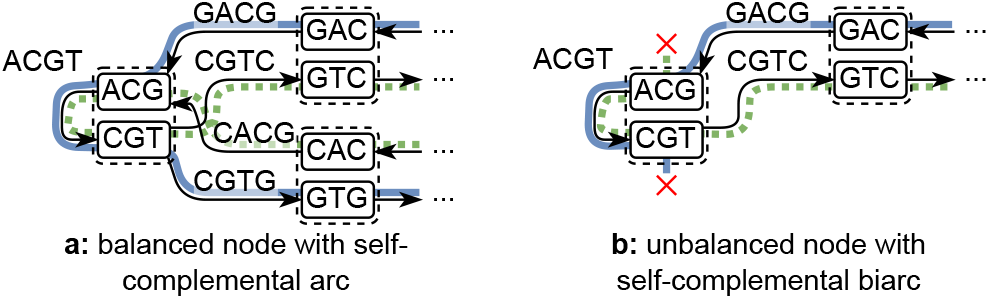
A self-complemental biarc [(*ACG, CGT*)] covered by a biEulerian circuit ([(*GAC, ACG*), (*CGT, GTC*)], [(*ACG, CGT*)], [(*CGT, GTG*), (*GAG, ACG*)]). The two directions of the bidirected circuit are drawn in blue ((*GAC, ACG*), (*ACG, CGT*), (*CGT, GTG*)) and green dotted ((*CAC, ACG*), (*ACG, CGT*), (*CGT, GTC*)). In (a), the binode [*ACG, CGT*] is balanced, hence the circuit can enter it with some biarc, cover the self-complemental biarc, and then leave it via some other biarc. If, like in (b), there was no other biarc to leave [*ACG, CGT*], then the graph would not be biEulerian, as the biarc [(*GAC, ACG*), (*CGT, GTC*)] cannot be used twice, even if the second use is in the other direction. Visually, the blue walk cannot use (*CGT, GTC*), since it was already used by the green dotted walk, and the green dotted walk cannot use (*GAC, ACG*), as it was already used by the blue walk.

#### Valid bidirected flow

To adapt Edmonds’ and Johnson’s formulation to biwalks, we additionally need to ensure that the resulting flow yields a set of biarcs, i.e. that each arc has the same flow as its reverse complement:

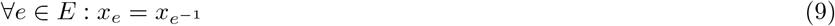

#### Adapted flow formulation

With the modifications above, we can adapt the formulation of Edmonds and Johnson to solve the bidirected partial-coverage Chinese postman problem. We define *F* to be the arcs in the original graph, and set *E*:= *V* × *V*. We further set *c_e_* = 1 for *e* ∈ *F* and *c_e_* = *k* – 1 otherwise. Lastly we define *S* and *T* as above. Then we get the following modified formulation:

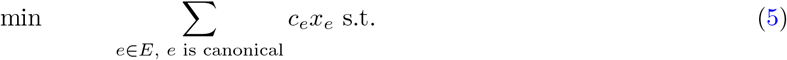

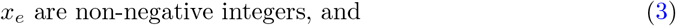

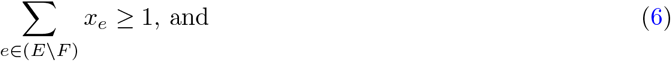

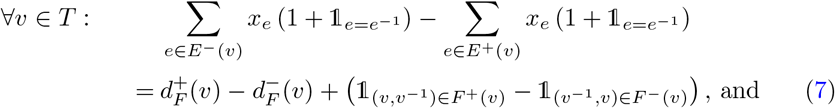

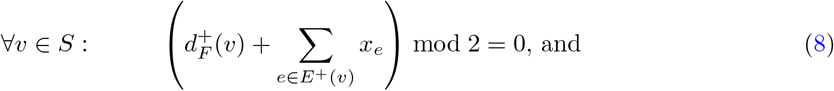

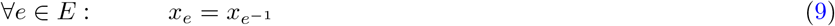

In this min-cost integer flow formulation^[12]^ of the bidirected partial-coverage Chinese postman problem, analogue to the formulation of Edmonds and Johnson, sources and sinks are nodes with missing outgoing or incoming arcs with demands matching the number of missing arcs in *F*. Our formulation would not be solvable for practical de-Bruijn graphs because inserting a quadratic amount of arcs into the graph is infeasible. However, most of the breaking arcs are not needed, since in a minimum solution they can only carry flow if they directly connect a source to a sink, by the following argument: Imagine a breaking arc that carries flow but is connected to a source or sink with at most one end. We can trace one unit of flow on the arc to a source and a sink, creating a path of flow one. By removing the flow from the path, and adding it to a breaking arc directly connecting the source to the sink, we get a valid flow. This flow has lower costs than the original, because it contains the same amount of breaking arcs, but a lower number of non-breaking arcs. This can be repeated until only breaking arcs that directly connect sources to sinks are left.

But even reducing the number of breaking arcs like this might not be enough if the graph contains too many sources and sinks. We therefore reduce the linear program to a min-cost matching instance, similar to Edmonds and Johnson.

### 5.6 Solving the min-cost integer flow formulation with min-cost matching

To solve the bidirected partial-coverage Chinese postman problem with min-cost matching, we observe that flow traverses the graph from a source to a sink only via min-cost paths, since all arcs have infinite capacity. Due to the existence of the breaking arcs with low cost (*k* – 1), we can further restrict the flow to use only paths of length at most *k* – 2 without affecting minimality. However, since we are also interested in a low number of strings in our minimum SPSS, we also allow paths of length *k* – 1. We can precompute these min-cost paths efficiently in parallel (see Section 5.7 below). Then it remains to decide which combination of min-cost paths and breaking arcs yield a minimum solution.

To simplify this problem, observe that the pairing of sources and sinks that are connected via breaking arcs does not matter. Any pairing uses the same amount of breaking arcs, and therefore has the same costs. It only matters that these nodes are not connected by a lower-cost path that does not use any breaking arcs, and that there is at least one breaking arc. As a result, we can ignore breaking arcs when searching a minimum solution, and instead introduce costs for unmatched nodes. However, we still need to enforce that there is at least one pair of unmatched nodes. We do this using a special construction described below. Note though, that there can only be unmatched nodes if there are unbalanced binodes, i.e. the graph was not biEulerian in the beginning. However, if the graph is biEulerian already, the whole matching step can be left out, and instead a biEulerian circuit with one arbitrarily inserted breaking biarc can be returned. So we can safely assume here that the graph contains at least one pair of unbalanced binodes^[13]^.

We formulate a min-cost matching problem with penalty costs for unmatched nodes, which can be reduced to a min-cost perfect matching problem. For the construction of our undirected matching graph *M* we define the set of sources *A* ⊆ *T* as all nodes with negative bi-imbalance, and the set of sinks *B* ⊆ *T* as all nodes with positive bi-imbalance. Then we add |*bi_v_*| (absolute value of the bi-imbalance of *v*) copies of each node from *A, B* and *S* to *M*. Further, for each min-cost path from a node *a* ∈ *A* ∪ *S* to a node *b* ∈ *B* ∪ *S* we add an edge (undirected arc) from each copy of a to each copy of *b* in *M* with costs equal to the costs of the path. We ignore self loops at nodes in *S* since they do not alter the imbalance, and nodes in *A* and *B* cannot have self loops.

Then, to fulfil Condition (9) (valid bidirected flow) and to reduce the size of the matching problem, we merge all nodes and arcs with their reverse complement (the unmerged graph is built here to simplify our explanations, in our implementation we directly build the merged graph). This additionally results in self-complemental biarcs forming self-loops in the merged graph, thus making them not chooseable by by the matcher. But this is correct, as self-complemental biarcs alter the bi-imbalance of a binode by two, and therefore they can only be chosen by matching two different copies of the same binode in *M*.

Additionally, to fulfil Condition (6) (cover one bidirected arc), we add a pair of nodes *u*, *w* to *M*. We connect *u* and *w* to each node in *M* (but do not add an edge between *u* and *w*) and assign costs 0 to all those edges. This forces *u* and *w* to be matched to other nodes *u*′, *w*′, which means that when biEulerising, the bi-imbalances of *u*′ and *w*′ need to be fixed with at least one breaking arc.

Lastly, we assign each node other than *u* and *w* penalty costs of (*k* – 1)/2 for staying unmatched, as each pair of unmatched nodes produces costs *k* – 1 for using a breaking arc.

We reduce *M* to an instance of the min-cost perfect matching problem using the reduction described in [64]. For that we duplicate the graph, and add an edge with costs *k* – 1^[14]^ between each node and its copy, but not between *v* and *w* and their respective copies.

After this reduction, we use Blossom V [65] to compute a solution. Since all nodes were doubled in the reduction, we actually get two solutions that might even differ, however both of them are minimum. We arbitrarily choose one of the solutions. This gives us a multiset of arcs that we complete with the breaking arcs required to balance the unmatched nodes to create a biEulerisation of the input graph. Following the approach from Edmonds and Johnson, we find a biEulerian circuit in the resulting graph which is a solution to the bidirected partial-coverage Chinese postman problem as required.

Note that our matching formulation only optimises CL, but does not optimise SC. It indirectly optimises SC because it decreases CL by joining strings, which also decreases SC by one each time. However, when joining two strings would not alter CL, Blossom V may output both variants, with joining the strings and without, while staying optimal. It then chooses an arbitrary option.

### 5.7 Efficient computation of many-to-many min-cost paths

Apart from solving the matching problem, finding the min-cost paths between sources and sinks is the most computationally heavy part of our algorithm.

We solve it using Dijkstra’s shortest path algorithm [66] in a one-to-many variant and execute it in parallel for all sources. To be efficient, we create a queue with blocks of source nodes, and the threads process one block at a time. A good block size balances between threads competing for access to the queue, and threads receiving a very imbalanced workload. Since our min-cost paths are short (at most *k* – 1 arcs), in most executions of Dijkstra’s algorithm only a tiny fraction of the nodes in the graph are visited. But the standard variant of Dijkstra’s algorithm wastefully allocates an array for all nodes to store their distance from the source node (the *distance array*). To save space, we instead use a hashmap, mapping from node_index to distance from source. This turned out to be faster than using a distance array, even if the distance array uses an epoch system^[15]^ to only do a full reset every 2^32^ queries. As another optimisation, we abort the execution early when Dijkstra reaches costs greater than *k* – 1, since we are only interested in paths up to costs *k* – 1.

Finally, in our implementation, we do not compute the actual sequences of arcs of the paths. Instead of copying the path arcs when biEulerising the graph, we insert special dummy arcs with a length equal to the length of the path. When breaking the final biEulerian circuit, if there are no breaking arcs but dummy arcs, then we break at a longest dummy arc to produce a minimum solution. If there are neither breaking nor dummy arcs, we proceed as described above. Then, when reporting the final set of strings, we define spell(·) to append the last *ℓ* characters of *b* when encountering a dummy biarc [(*a, b*), (*b*^−1^, *a*^−1^)] (or [(*a, b*)]) of length *ℓ*.

### 5.8 Efficient computation of the greedy heuristic

The greedy heuristic biEulerises the graph by greedily adding min-cost paths between unbalanced nodes, as opposed to choosing an optimal set via min-cost matching like our main algorithm. It then continues like the main algorithm, finding a biEulerian circuit, breaking it into walks and spelling out the strings.

To be efficient, the min-cost paths are again computed in parallel, and we apply all optimisations from Section 5.7. The parallelism however poses a problem for the greedy computation: if a binode with one missing incoming biarc is reached by two min-cost paths in parallel, then if both threads add their respective biarcs, we would overshoot the bi-imbalance of that binode. To prevent that, we introduce a lock for each node, and before inserting a biarc into the graph we lock all (up to) four incident nodes. By locking the nodes in order of their ids we ensure that no deadlock can occur. Since the number of threads is many orders of magnitude lower than the number of nodes, we assume that threads almost never need to wait for each other. In addition to the parallelisation, we abort Dijkstra’s algorithm early when we have enough paths to fix the imbalance for the binode. This sometimes requires to execute Dijkstra’s algorithm again if a potential sink node was used by a different thread in parallel. But again, since the number of threads is many orders of magnitude lower than the number of nodes, we assume that this case almost never occurs.

In practice, the greedy heuristic achieves better results in terms of SC than the optimal matchtigs algorithm (see the Results sections). This is because the greedy heuristics always joins two strings if it does not alter CL, while the optimal algorithm does not, as explained in Section 5.6.

### 5.9 Minimising string count

In the paper we studied SPSSes of minimum total length (minimum CL). In this section we note that an SPSS with a minimum number of strings (minimum SC), and with no constraint on the total length, is also computable in polynomial time.

The high-level idea, ignoring reverse complements, is as follows. Given the arc-centric de Bruijn graph *G*, construct the directed acyclic graph *G*\* of strongly connected components (SCCs) of *G*. In *G*\*, every SCC is a node, and we have as many arcs between two SCCs as there are pairs of nodes in the two SCCs with an arc between them. Clearly, all arcs in a single SCC are coverable by a single walk. Moreover, for two SCCs connected by an arc, their two covering walks can be connected via this arc into a single walk covering all arcs of both SCCs. Thus, the minimum number of walks needed to cover all arcs of *G*\* (i.e., minimum SC SPSS) equals the minimum number of paths needed to cover all arcs of the acyclic graph *G*\*. This is a classic problem solvable in polynomial time with network flows (see e.g. [67] among many).

However, such an SPSS of minimum SC very likely has a large CL, because covering an SCC with a single walk might repeat quadratically many arcs, and connecting the covering walks of two adjacent SCCs might additionally require to repeat many arcs to reach the arc between them.

### 5.10 Experimental evaluation

We ran our experiments on a server running Linux with two 64-core AMD EPYC 7H12 processors with 2 logical cores per physical core, 1.96TiB RAM and an SSD. We downloaded the genomes of the model organisms from RefSeq [42]: Caenorhabditis elegans with accession GCF_000002985.6, Bombyx mori with accession GCF_000151625.1 and Homo sapiens with accession GCF_000001405.39. These are the same genomes as in [43], except that we downloaded HG38 from RefSeq for citability. The short reads were downloaded from the sequence read archive [68]: Caenorhabditis elegans with accession SRR14447868.1, Bombyx mori with accession DRR064025.1 and Homo sapiens with accessions SRR2052337.1 to SRR2052425.1.

For the pangenomes we downloaded the 616 Streptococcus pneumoniae genomes from the sequence read archive, using the accession codes provided in Table 1 in [69]. We downloaded the 1102 Neisseria gonorrhoeae genomes from [70]. Up to here the pangenomes are retrieved in the same way as in [43]. We additionally used `grep` to select 3682 Escherichia coli genomes from ftp://ftp.ncbi.nlm.nih.gov/genomes/genbank/bacteria/assembly_summary.txt. The ~309k salmonella genome sequences were downloaded from the EnteroBase database [71], gzippedí^16]^. The 2505x human pangenome is from the 1000 genomes project [72], created by downloading a variant of GRCh37 from ftp://ftp.1000genomes.ebi.ac.uk/vol1/ftp/technical/reference/phase2_reference_assembly_sequence/hs37d5. fa.gz and downloading variant files for chromosomes 1-22 from http://ftp.1000genomes.ebi.ac.uk/vol1/ftp/release/20130502/. We then converted chromosomes 1-22 in the reference into 2505 sequences each using the tool `vcf2multialign` published in [73].

For querying the human read dataset, we used 3 million reads randomly drawn from the reads used to construct the dataset. For querying the Salmonella pangenome, we used 3 million randomly drawn short reads from 10 read data sets from the sequence read archive with the accessions listed in Additional file 9. For querying the human pangenome, we used 3 million randomly drawn short reads from one file of the sequence read archive with accession SRR2052337.1. This is one of the files from the human short read dataset described above. For querying the E.coli pangenome (in Additional file 6) we used 30 short read data sets from the sequence read archive with the accessions listed in Additional file 7.

We used snakemake [74] and the bioconda software repository [75] to craft our experiment pipeline. The tigs were checked for correctness by checking the kmer sets against unitigs. The bifrost queries in Additional file 6 were checked for correctness by checking that the query results are equivalent for those with unitigs. The SSHash queries were not checked for correctness, as SSHash was modified by the author himself. Whenever we measured runtime of queries and builds for Additional file 5 (Performance with different amounts of threads), we only let a single experiment run, even if the experiment used only one core. When running the other builds we ran multiple processes at the same time, but never using more threads than the processor has physical cores (thus avoiding any effects introduced by sharing logical cores). When running a tool we copied its input to the SSD, and copied the results back to our main network storage, to prevent the network storage’s varying workload to affect our runtime measurements. For experiments running on input reads or references as opposed to unitigs (BCALM2, ProphAsm), we copied the inputs to a RAID of HDDs instead, due to their large size. The copying was not part of the measurements. We made sure that the server is undisturbed, except that we monitored the experiment status and progress with `htop` and `less`. We limited each run to 256GiB of RAM per process, which prevented us from running matchtigs on larger inputs. Further, ProphAsm supports only *k* ≤ 32, so it was not run for *k* larger than that.

For an overview of our experiment pipeline for computing tigs, see Figure 5. We run ProphAsm on the input data, as it was introduced to do [43]. All other tools require unitigs to be computed first. UST specifically requires unitigs computed by BCALM2, as BCALM2 adds additional annotations to the fasta unitig file. Our tool matchtigs also can make use of these annotations to speed up the construction of the arc-centric de Bruijn graph. On the human pangenome, BCALM2 crashed due to the input being too large. Hence we used Cuttlefish 2 [40] to compute unitigs, and since UST only runs on unitigs computed by BCALM2, we then ran ProphAsm to compute heuristic simplitigs.

**Figure 5:**
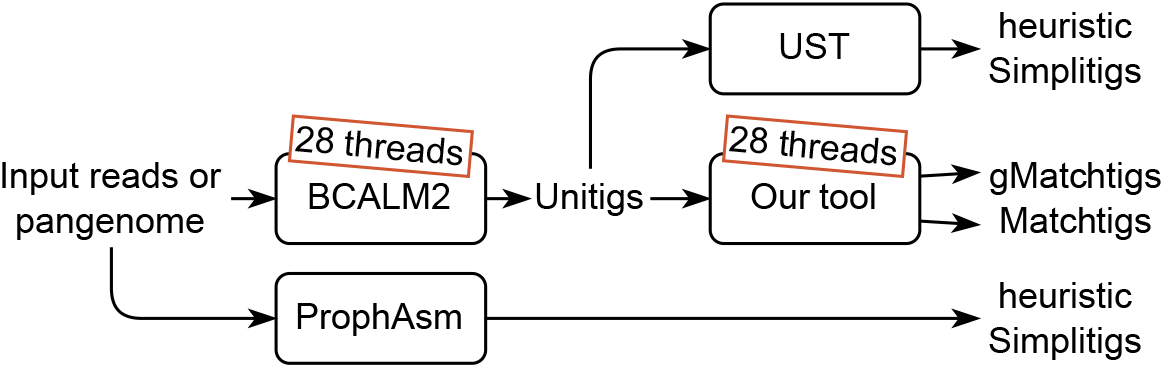
The DAG of tools run on the short read sets and pangenomes to compute different tigs.

For queries, we executed Bifrost or SSHash-Lite on the different tigs. The Bifrost query command handles both building the index and executing the query, while SSHash-Lite requires to run a separate command to build the index first.

See Section *Availability of data and materials* for availability of our implementation and experiment code, which includes all the concrete commands we have used to execute our experiments.

## Acknowledgements

We are very grateful to Paul Medvedev and Amatur Rahman for helpful initial discussions on this problem. We further wish to thank the Finnish Computing Competence Infrastructure (FCCI) for supporting this project with computational and data storage resources. We also wish to thank Andrea Cracco for providing us with the ~309kx Salmonella pangenome. We further wish to thank the anonymous reviewers for there useful constructive feedback, which improved the presentation of the paper, the implementation and the experimental results. Finally we wish to thank the Rust community (https://users.rust-lang.org) for explanations about language-specific details of parallel implementations.

## Funding

This work was partially funded by the European Research Council (ERC) under the European Union’s Horizon 2020 research and innovation programme (grant agreement No. 851093, SAFEBIO), and by the Academy of Finland (grants No. 322595, 328877). This work was also partially supported by the project MobiDataLab (EU H2020 RIA, grant agreement N-101006879).

## Availability of data and materials

The implementation of the matchtigs and greedy matchtigs algorithms is available on github at https://github.com/algbio/matchtigs. The name of the project is *matchtigs*. It is platform independent, and can be compiled locally or installed from bioconda as described in the README of the project. It is licensed under the 2-clause BSD license. The version used for our experiments is available at https://doi.org/10.5281/zenodo.7275977, and the implementation together with all code to reproduce the experiments is available at https://doi.org/10.5281/zenodo.7275990. The experiment code is also licensed under the 2-clause BSD license. SSHash-Lite is available at https://doi.org/10.5281/zenodo.7277145 and licensed under the MIT license. See Section 5.10 for the availability of the non-original data used for our experiments.

## Competing interests

The authors declare that they have no competing interests.

## Authors’ contributions

SK and AIT formulated the problem. AIT and SK designed an optimal algorithm and SK implemented and evaluated its prototype. Thereafter, SS improved the algorithm’s design to its current form and provided the final implementation as well as optimisations required to make it practically relevant. SS developed the greedy heuristic and implemented it. JA, SS and AIT designed the experiments and interpreted the results. SS performed all experiments, and developed all further code published in the context of this work. SS wrote the manuscript under the supervision of the other authors, except for Section 2.5. GEP created and described SSHash-Lite in Section 2.5 and ran the query experiments on the machine with focus on single-core performance. All authors reviewed and approved the final version of the manuscript.

## Additional Files

Additional file 1 — Quality of compressing model organisms (pdf)

Additional CL and SC data from the experiments from Table 1, with varying *k* and min. abundance. Also, the average unitig length and total unitig count is plotted.

Additional file 2 — Performance of compressing model organisms (pdf)

Additional run time and memory data from the experiments from Table 1, with varying k and min. abundance.

Additional file 3 — Quality of compressing pangenomes (pdf)

Additional CL and SC data from the experiments from Table 2, with varying k and min. abundance. Also, the average unitig length and total unitig count is plotted.

Additional file 4 — Performance of compressing pangenomes (pdf)

Additional run time and memory data from the experiments from Table 2, with varying k and min. abundance.

Additional file 5 — Performance with different amounts of threads (pdf)

A run time and memory comparison of compressing the E. coli pangenome and the H. sapiens reference genome with different amounts of threads and different compression methods.

Additional file 6 — Query experiment on E.coli with Bifrost (pdf)

The query experiment of Section 2.5 using Bifrost as query tool and querying on E.coli.

Additional file 7 — Accessions of the reads used for the E.coli query experiment (txt)

A list of sequence read archive accession numbers for the 30 sets of E.coli short reads used for querying.

Additional file 8 — Proofs of optimality (pdf)

Proofs for the optimality of our algorithm.

Additional file 9 — Accessions of the reads used for the Salmonella query experiment (txt)

A list of sequence read archive accession numbers for the 10 sets of Salmonella short reads used for querying.

Additional file 10 — Query experiment on a machine with focus on single-core performance (pdf) The query experiment of Section 2.5 on a machine with focus on single-core performance.

## Footnotes

[1] Our own observation.

[2] In [44], an SPSS is defined for a given set of kmers as a set of strings that contains the same spectrum of kmers, where the spectrum is defined as a multiset. Since a set of kmers contains each kmer at most once, this implies that kmers must be unique in such a definition of an SPSS. However, the spectrum is usually defined as a set and not a multiset, hence we use the term SPSS to describe a set of strings that contains the same set of kmers as the input, allowing repetitions.

[3] There might even be cases where by increasing CL and decreasing SC, the overall size of the representation of the string set (strings + index structure) can be decreased. To stay independent of any specific data structure we only optimise CL.

[4] While this paper was under review, Schmidt and Alanko realised that the algorithm to compute matchtigs can also be used to compute optimal simplitigs, by leaving out all parts related to repeating kmers. The matchtig algorithm can hence be seen as an extension to the Eulertig algorithm, even if the former was discovered first.

[5] Personal communication by Andrea Cracco.

[6] ProphAsm supports only *k* ≤ 32 such that a comparison is often impossible. But where it is possible, it performs only slightly better than UST in terms of CL and SC.

[7] https://github.com/algbio/matchtigs

[8] The code is available at https://github.com/jermp/sshash-lite.

[9] This is because there may be multiple circular original-biarc-covering biwalks with minimum costs, but with different repeated kmers. When breaking the walk by removing a longer sequence of repeated kmers, the resulting string gets shorter, the more repeated kmers get removed.

[10] Either by connecting nodes with missing outgoing arcs directly to nodes with missing incoming arcs, or by connecting them via a path of multiple arcs.

[11] Since the labels of an arc are of length_*k* – 1_ and those of a node are of length k-1, only one can have even length, so only one can be self-complemental for DNA alphabets.

[12] It is conceptually similar to that proposed in [56], however different because the basic definitions differ, and we further allow for special arcs of which only one needs to be covered.

[13] It cannot contain a single unbalanced binode, see e.g. [47]

[14] By duplicating the graph, virtually each edge’s costs are doubled, since each edge exists twice afterwards. However, the edges between nodes and their duplicate exist only once, so their costs require doubling.

[15] An epoch system stores a second value for each entry in the distance array indicating in what execution of Dijkstra’s algorithm that value is valid. The execution counter gets incremented each execution, and only when it wraps around, the distance array is reset normally. Whenever an entry in the distance array is read, the value infinity is returned if the execution counter does not match the execution saved for the entry in the distance array. Whenever an entry in the distance array is written, the corresponding execution value is set to the current value of the execution counter.

[16] Downloaded in February 2022.

## Notes

https://github.com/algbio/matchtigs

https://github.com/jermp/sshash-lite

### Competing Interest Statement

The authors have declared no competing interest.

### Summary of Updates

Forgot to update abstract

