## Additional file 1 for "Matchtigs: minimum plain text representation of kmer sets"

### Additional file 1: Quality of compressing model organisms

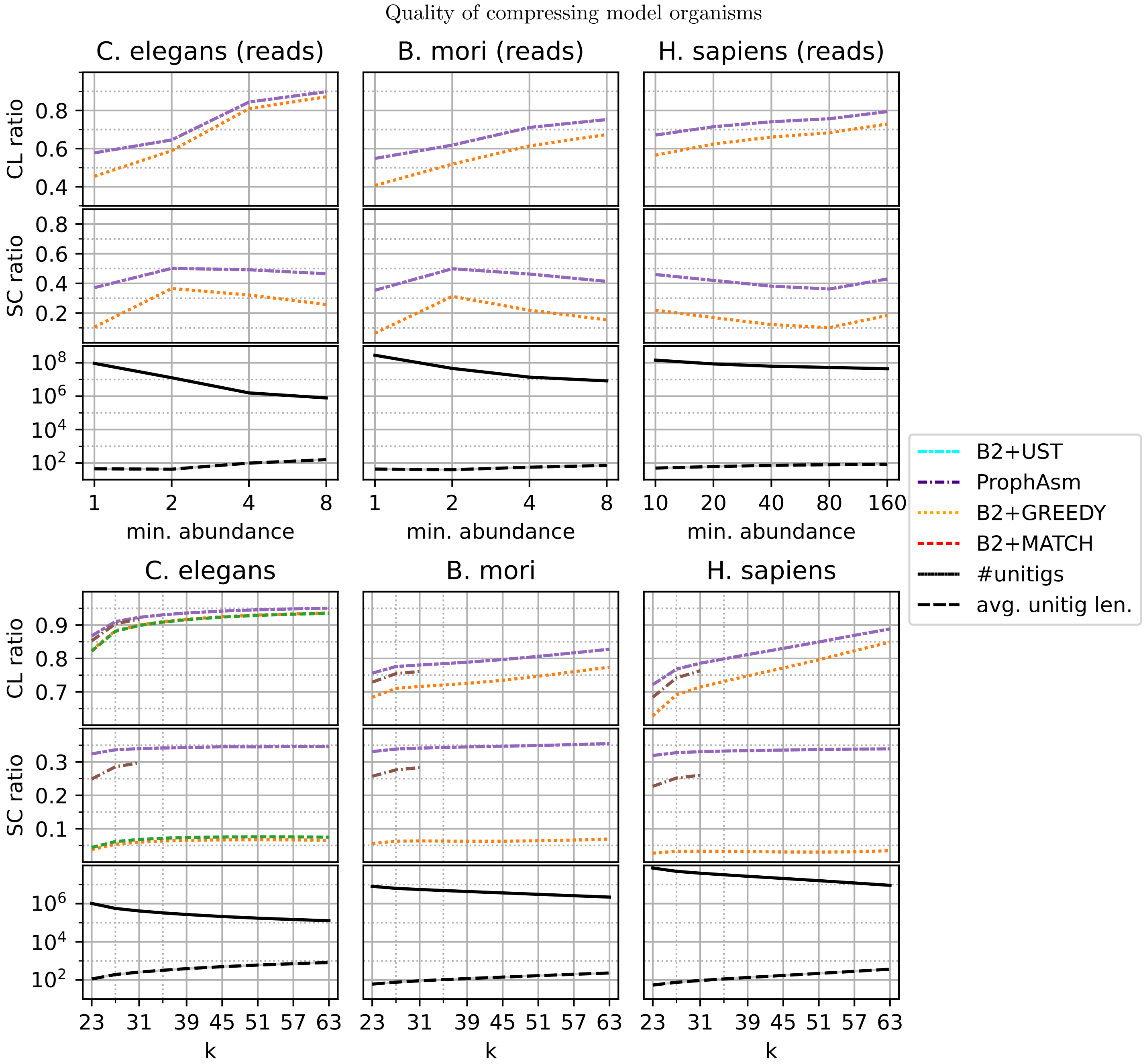


Figure 1: Ratio of CL and SC between different compression methods and unitigs, as well as the number and average length of unitigs for reference genomes and short reads of model organisms. For read data sets we chose $k = 51$, and for the reference genomes we chose a min abundance of 1. ProphAsm handles only $k \leq 32$ and matchtigs are memory-feasible only for C. elegans, so only the corresponding runs are shown. ProphAsm and UST produce overlapping lines in all subplots, and matchtigs and greedy matchtigs mostly overlap. The lengths of the genomes are 100Mbp for Caenorhabditis elegans, 482Mbp for Bombyx mori and 3.21Gbp for Homo sapiens and the read data sets have a coverage of 64x for Caenorhabditis elegans, 58x for Bombyx mori and 300x for Homo sapiens.
