## Additional file 3 for "Matchtigs: minimum plain text representation of kmer sets"

### Additional file 3: Quality of compressing pangenomes

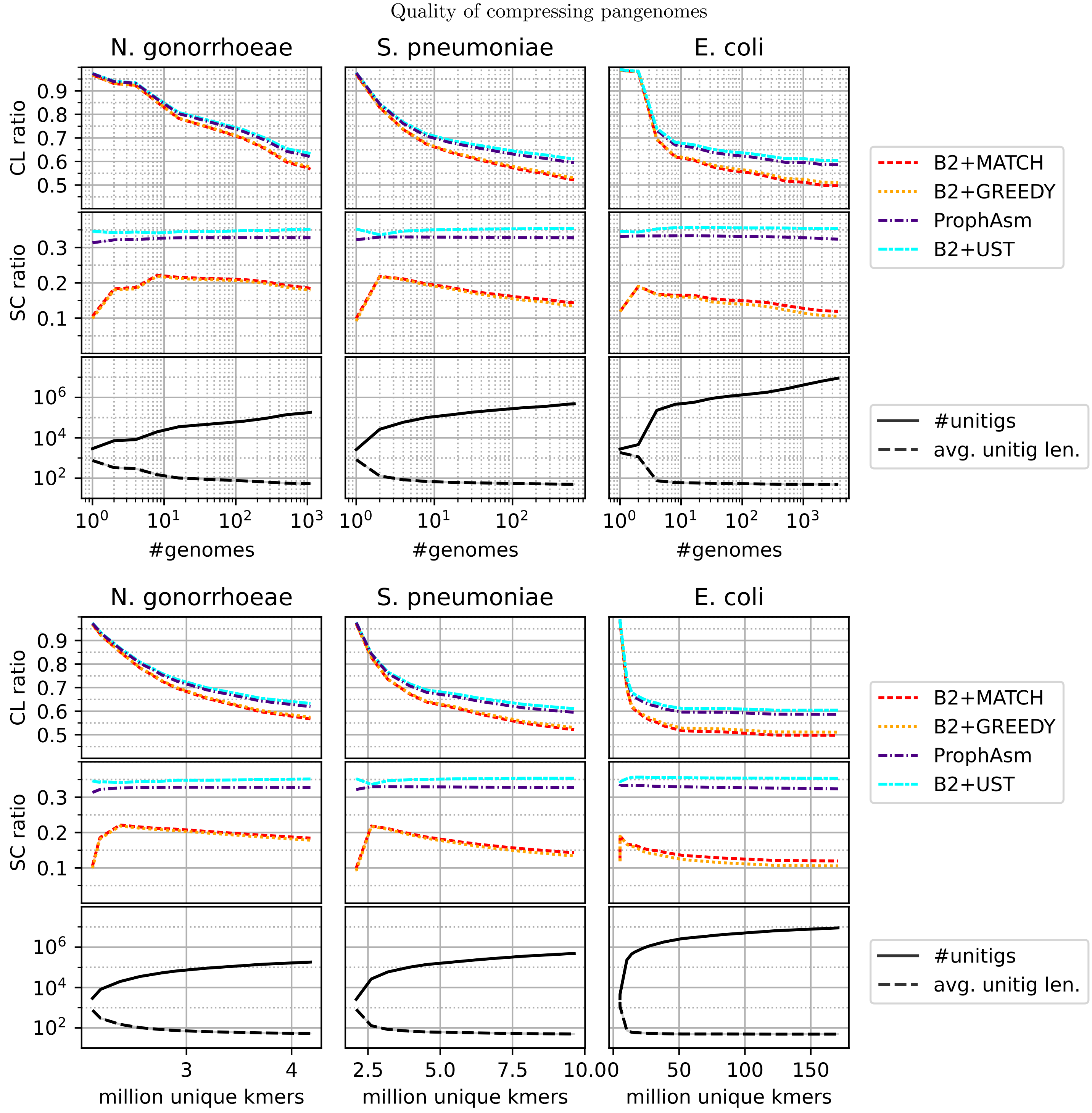


Figure 1: Ratio of CL and SC between different compression methods and unitigs, as well as the number and average length of unitigs for pangenomes. We chose $k = 31$, and a min abundance of 1. ProphAsm and UST produce overlapping lines in all subplots, and matchtigs and greedy matchtigs mostly overlap. The lengths of the genomes are 2.15Mbp for Neisseria gonorrhoeae, 2.22Mbp for Streptococcus pneumoniae and 4.64Mbp for Escherichia coli.
