## Additional file 4 for "Matchtigs: minimum plain text representation of kmer sets"

### Additional file 4: Performance of compressing pangenomes

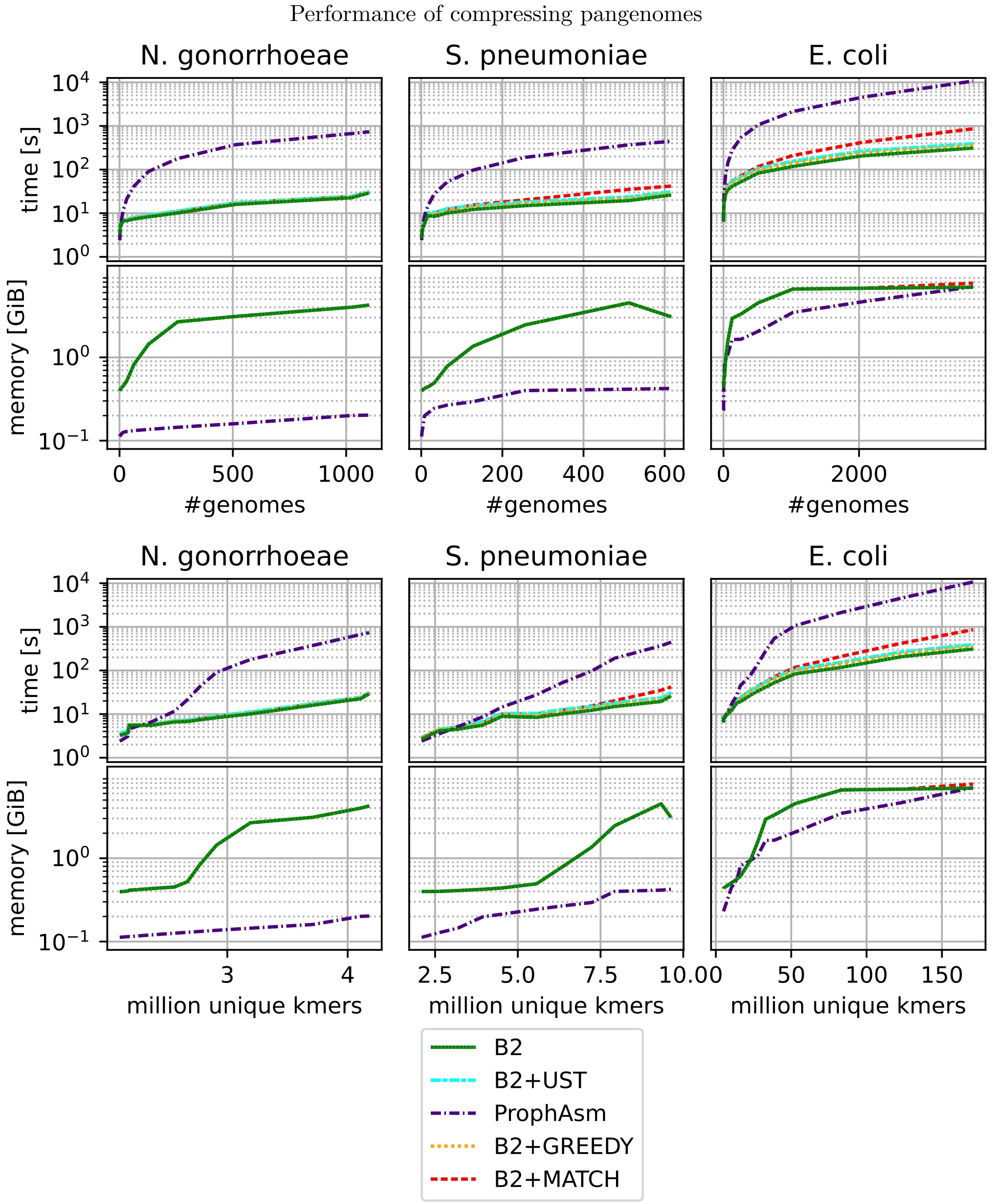


Figure 1: Time and memory consumption of different compression methods for pangenomes. We chose $k = 31$ and a min abundance of 1. The time and memory of algorithms including BCALM2 overlap often, since they are dominated by BCALM2. BCALM2, greedy matchtigs and matchtigs are run with 28 threads, while UST and ProphAsm can only be run with one thread (for UST, the preceding run of BCALM2 was executed with 28 threads). For UST, greedy matchtigs and matchtigs, note that they take unitigs as input, so they require a run of BCALM2 as preprocessing. The time in these cases is given as the sum of the time taken by both tools, and the memory as the maximum.
