## Additional file 5 for "Matchtigs: minimum plain text representation of kmer sets"

### Additional file 5: Performance with different amounts of threads

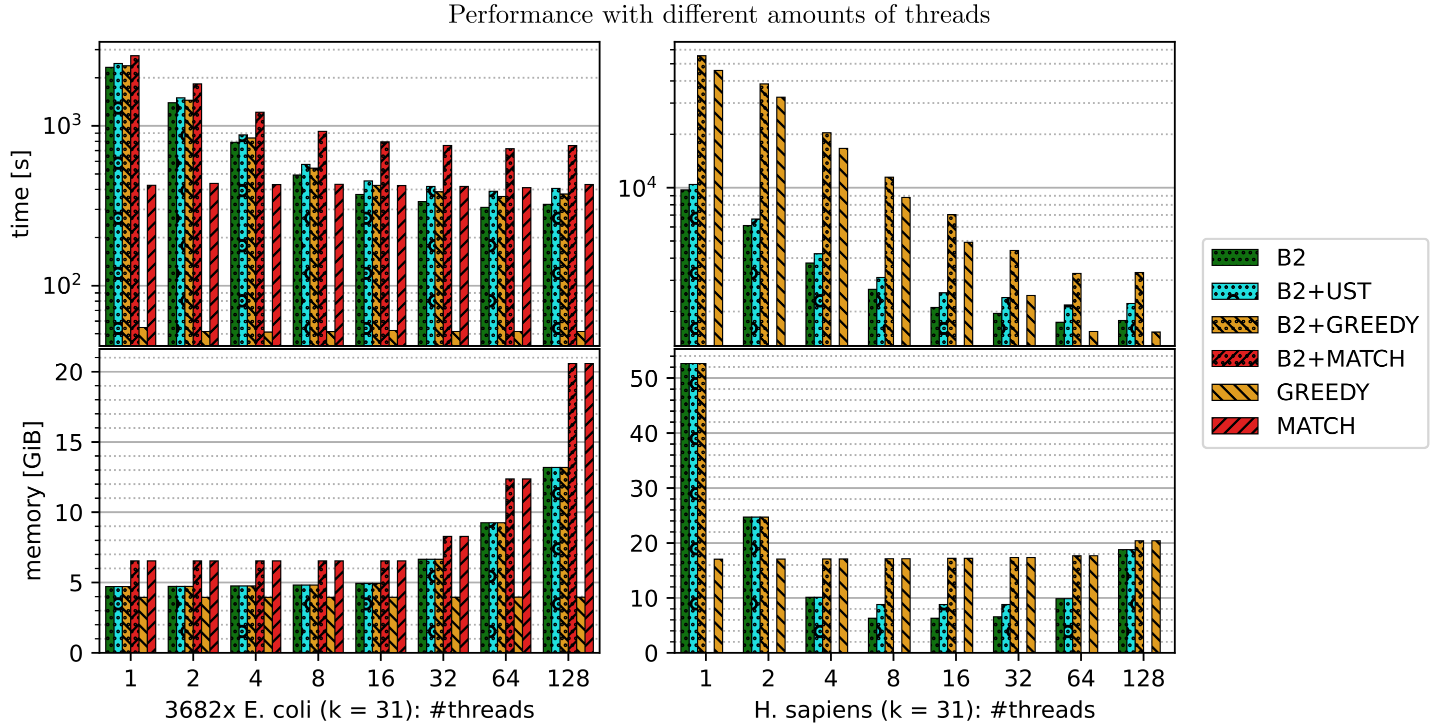


Figure 1: Time and memory consumption of different compression methods for the Escherichia coli pangenome and the Homo sapiens genome. We chose a min abundance of 1 and varied the number of threads. Since UST cannot be run with more than one thread, it is always run with one thread (the preceding run of BCALM2 was executed with the correct number of threads). For UST, greedy matchtigs and matchtigs, note that they take unitigs as input, so they require a run of BCALM2 as preprocessing. The time in these cases is given as the sum of the time taken by both tools, and the memory as the maximum.
