## Additional file 6 for "Matchtigs: minimum plain text representation of kmer sets"

### Additional file 6: Query experiment on E.coli with Bifrost

Table 1: Performance characteristics of querying different tigs with Bifrost.

| genome | algorithm | indexing [s] | searching [s] | total [s] | search speedup | | mem [GiB] |
|---|---|---|---|---|---|---|---|
| 3682x E. coli | unitigs | 21.00 | 336.79 | 360.37 | 1.00 | | 2.39 |
| | UST | 16.11 | 267.02 | 285.52 | 1.26 | | 2.13 |
| | ProphAsm | 15.55 | 261.65 | 278.91 | 1.29 | | 2.10 |
| | gMatchtigs | 16.19 | 208.10 | 226.23 | 1.62 | (1.26) | 2.12 |
| | matchtigs | 16.33 | 202.71 | 220.73 | 1.66 | (1.29) | 2.09 |

The Bifrost query command is run with 16 threads and default settings. 3682 E.coli genomes are queried with all raw reads used to assemble 30 of the genomes. All columns are averages of 10 experiments. The search speedup is with respect to unitigs, and the search speedup in parentheses is with respect to the strings computed by ProphAsm. Indexing is the time required to build the minimizer index within Bifrost, and searching is the time required to check if a read is in the pangenome using the minimizer index. The total time is the wall-clock time of bifrost.

Matchtigs have further applications beyond merely reducing the size required to store a set of kmers. Due to their smaller size and lower string count, they can make downstream applications more efficient. For example, the kmer-based query tool Bifrost [1] achieves speedups of 1.66 when using matchtigs instead of unitigs, and 1.29 when using matchtigs instead of the strings computed by ProphAsm (see Table 1 for the detailed results). Note that we measure the speedup of the search phase only, since the index phase is independent of the query and therefore could be moved into a separate preprocessing step. Note also that these speedups are consistent with those reported when using heuristic simplitigs [2] for kmer queries with BWA-MEM [3] (without modification of BWA-MEM).

We achieve our speedups *without modifying Bifrost*[1], but just by passing Bifrost a file with our tigs instead of unitigs. When using in query mode, Bifrost is intended to take a GFA file containing a compacted de Bruijn graph as well as a file containing the query strings as inputs. It then builds a minimizer index over the unitigs in the GFA file, and executes the queries using this index. If we pass a GFA file containing e.g. matchtigs instead of a compacted de Bruijn graph, then Bifrost still works the same way. It builds a minimizer index over the matchtigs inside the GFA file, and then uses that index to execute the queries. It does not build a compacted de Bruijn graph from the matchtigs, or in any other way use any topology information that would be in the GFA file. This is why the query still works correctly, even though the GFA file does not contain a compacted de Bruijn graph.

The speedups that we achieve when passing e.g. matchtigs instead of unitigs in the GFA file can be explained as follows. When querying, Bifrost iterates over the kmers of the query. It finds the tig containing the kmer using the minimizer index and then extends that match until the query and the tig differ, or the query or tig ends. In that case, it again queries the minimizer index, until all query kmers are checked. When using longer tigs with a lower tig count, then the number of queries to the minimizer index is lower, as the end of a tig is reached more rarely. Extending a match is extremely fast, as it only requires a comparison between the character following the last match on the tig and the character following the last match on the query. However querying the minimizer index requires to compute the hash of the current minimizer, as well as looking up the hash inside the hash table, and then checking which tig does actually contain the original kmer, and not just the minimizer. Therefore, by reducing the number of queries to the minimizer index, we can reduce the overall runtime of the query.

In addition to speedups, a smaller representation is also expected to decrease the memory consumption. However in practice the memory consumption only increases when switching from unitigs to heuristic simplitigs, but not when further switching to matchtigs or greedy matchtigs. We assume that this is due to the longer strings requiring more memory while being loaded block-wise in ASCII format before they get stored in a two-bits-per-character compressed format.

[1] We did need to add code to measure index and search time separately.
