## Additional file 8 for "Matchtigs: minimum plain text representation of kmer sets"

### Additional file 8: Proofs of optimality

In this subsection we provide proof for the minimality of the SPSS computed by our algorithm. We call the input set of strings $I$, and the output SPSS $S$. We assume that the graph $DBG_k(I)$ is connected. If it is not, then executing our algorithm on each maximal connected subgraph yields an optimal solution. This is because there can only be breaking arcs between different maximal connected subgraphs, i.e. every string in $S$ is within only a single maximal connected subgraph. Therefore, processing the maximal connected subgraphs separately does not decrease the solution space.

We recite some definitions from the main matter for convenience.

**Definition 1** (Graph transformation)**.** *Given an arc-centric de-Bruijn graph $DBG_k(I) = (V, E)$, the* transformed graph *is defined as $DBG'_k(I) = (V, E')$ where $E'$ is a multiset defined as $E' := E \cup (V \times V)$. In $E'$, arcs from $E$ are marked as* non-breaking*, and arcs from $V \times V$ are marked as* breaking *arcs. The* cost function *$c(e), e \in E'$ assigns all non-breaking arcs the costs $1$ and all breaking arcs the costs $k - 1$.*

**Definition 2** (Circular original-biarc-covering biwalk)**.** *Given a transformed graph $DBG'_k(I) = (V, E)$, a* circular original-biarc-covering biwalk *is a circular biwalk $w$ such that for each non-breaking arc $(a, b) \in E$ there is a biarc $[(a, b), (b^{-1}, a^{-1})]$, $[(b^{-1}, a^{-1}), (a, b)]$ or $[(a, b)]$ in $w$. Additionally, $w$ needs to contain at least one breaking biarc.*

**Definition 3** (Costs of a circular original-biarc-covering biwalk)**.** *Given a transformed graph $DBG'_k(I)$ and a circular original-biarc-covering walk $w$ possibly consisting of biarcs $[(a, b), (b^{-1}, a^{-1})]$ and self-complemental biarcs $[(a, b)]$. The* costs *$c(w)$ of $w$ are the sum of the costs of each biarc and self-complemental biarc, where the costs of a biarc $[(a, b), (b^{-1}, a^{-1})]$ are $c((a, b))$, and the costs of a self-complemental biarc $[(a, b)]$ are $c((a, b))$.*

#### Reduction to the bidirected partial-coverage Chinese postman problem.

First, we show that the reduction to the bidirected partial-coverage Chinese postman problem is correct. For that we assume that we are given a solver for this variant of the Chinese postman problem, and formally recite our algorithm as follows:

**Algorithm 4** (Chinese-postman-based algorithm)**.**

1. *Compute the arc-centric de Bruijn graph $DBG_k(I)$.*
2. *Compute the transformed graph $DBG'_k(I)$ (Definition 1).*
3. *Use the given solver to compute a circular original-biarc-covering biwalk $w^*$ of minimum cost (Definition 2).*
4. *Break $w^*$ at all breaking biarcs.*
5. *Output set of strings $S$ spelled by the resulting walks.*

We first show that the algorithm only outputs SPSS and that the size of the final SPSS is equal to the costs of the biwalk $w^*$ computed in the algorithm. Using these lemmas, we can finally prove that the SPSS output by the algorithm is minimum.

**Lemma 5** (The Chinese-postman-based algorithm outputs an SPSS)**.** *When applying Steps 4 and 5 of Algorithm 4 to a circular original-biarc-covering biwalk $w$ in a transformed graph $DBG'_k(I)$, then the result is an SPSS of $I$.*

*Proof.* Observe that $w$ contains each non-breaking biarc of $DBG'_k(I)$ by definition. When being broken in Step 4, only breaking biarcs are removed and hence only biarcs that are not connected to any input kmer. So, the broken biwalks contain all biarcs of $DBG_k(I)$. Furthermore, all breaking biarcs are removed, and the remaining biarcs all stem from $DBG_k(I)$. So, the broken biwalks contain only biarcs of $DBG_k(I)$, and therefore the same set of biarcs as $DBG_k(I)$. Therefore, the strings spelled in step 5 of Algorithm 4 contain the same kmers as the input (but they may be reverse complemented), and hence have the same spectrum. Concluding, $S$ is an SPSS of $I$. □

**Lemma 6** (Costs equal size)**.** *When applying Steps 4 and 5 of Algorithm 4 to a circular original-biarc-covering biwalk $w$ in a transformed graph $DBG'_k(I)$, the resulting SPSS $S$ has a size equal to the costs of $w$.*

*Proof.* We denote the result of breaking $w$ at all breaking biarcs as $W$. Each walk $w' \in W$ spells a string $s' \in S$ and it holds that $|w'| + (k - 1) = |s'|$. Further, $w$ contains a breaking biarc for each walk in $W$, and we have assigned costs $k - 1$ to each breaking biarc, and costs of 1 to each other biarc. Therefore, the costs of $w$ equal the size of $S$. □

**Lemma 7** (Optimality of the Chinese-postman-based algorithm)**.** *The set of strings $S$ output by Algorithm 4 is a minimum SPSS of $I$.*

*Proof.* Algorithm 4 computes a minimum-cost circular original-biarc-covering biwalk $w^*$ in the transformed graph $DBG'_k(I)$. It computes a set of strings $S$ from $w^*$ by applying steps 4 and 5 where $S$ is an SPSS of $I$ by Lemma 5. By Lemma 6, the costs of any minimum-cost circular original-biarc-covering biwalk $w'$ are equal to the size of the SPSS $S'$ that is produced by applying Steps 4 and 5 to $w'$. So, since each SPSS can be represented by some $w'$, the existence of an SPSS smaller than $S$ would imply the existence of a minimum-cost circular original-biarc-covering biwalk cheaper than $w^*$, which contradicts the minimality of $w^*$. Therefore, $S$ is of minimum size. □

#### Solving the bidirected partial-coverage Chinese postman problem with min-cost integer flows.

Given that our algorithm is correct if we feed it a correct minimum-cost circular original-biarc-covering biwalk, we need to show that we actually do. For that, we start by showing that a solution to the flow formulation in Section 5.5 correctly results in a minimum biEulerisation which results in a minimum-cost circular original-biarc-covering biwalk as required. We restate the flow formulation and all defined variables for the reader's convenience:

**Definition 8** (Flow formulation)**.** *Given a transformed graph $DBG'_k(I) = (V, E)$, we define*

- *$F$ to be the non-breaking arcs in $E$,*
- *$c_e := 1$ for $e \in F$ and $c_e := k - 1$ for $e \in (E \setminus F)$, and*
- *$S$ to be the set of all self-complemental nodes and $T := (V \setminus S)$.*

*Then the* flow formulation *of the bidirected partial-coverage Chinese postman problem in $DBG'_k(I)$ is:*

$$\min \sum_{e\in E,\ e \text{ is canonical}} c_e x_e \text{ s.t.} \tag{1}$$

$$x_e \text{ are non-negative integers, and} \tag{2}$$

$$\sum_{e\in(E\setminus F)} x_e \geq 1, \text{ and} \tag{3}$$

$$\forall v \in T: \quad \sum_{e\in E^-(v)} x_e\left(1 + \mathbb{1}_{e=e^{-1}}\right) - \sum_{e\in E^+(v)} x_e\left(1 + \mathbb{1}_{e=e^{-1}}\right) = d_F^+(v) - d_F^-(v) + \left(\mathbb{1}_{(v,v^{-1})\in F^+(v)} - \mathbb{1}_{(v^{-1},v)\in F^-(v)}\right), \text{ and} \tag{4}$$

$$\forall v \in S: \quad \left(d_F^+(v) + \sum_{e\in E^+(v)} x_e\right) \bmod 2 = 0, \text{ and} \tag{5}$$

$$\forall e \in E: \quad x_e = x_{e^{-1}} \tag{6}$$

Additionally, we formalise this part of our algorithm:

**Algorithm 9.**

1. *Compute the arc-centric de Bruijn graph $DBG_k(I)$.*
2. *Compute the transformed graph $DBG'_k(I)$ (Definition 1).*
3. *Build and solve the flow problem according to Definition 8.*
4. *Insert $x_e$ copies of each arc $e \in E$ with flow value $x_e \geq 1$, and set the multiplicity of breaking arcs $e$ to $x_e$ (removing breaking arcs $e$ with a flow value $x_e = 0$).*
5. *Compute a biEulerian circuit (see e.g. [1] for details).*

The final biEulerian circuit is the minimum-cost circular original-biarc-covering biwalk of $DBG'_k(I)$. To show that this is true, in the same pattern as above we first show that the algorithm only outputs valid circular original-biarc-covering biwalks, and then show that the costs that it optimises are equivalent to the costs of the output walk.

we first prove that the costs optimised by the flow problem can be transformed into the costs in the Chinese postman problem by addition of a constant. Then we show that for each (minimum) solution to the Chinese postman problem $w$, there exists a solution $x$ to the flow instance that produces a solution $w'$ to the Chinese postman problem if fed into steps 4 and 5 of Algorithm 9, where the costs $c(w)$ of $w$ are equal to the costs $c(w')$ of $w'$. This is enough to argue that Algorithm 9 produces a minimum-cost circular original-biarc-covering biwalk of $DBG'_k(I)$.

**Lemma 10.** *Algorithm 9 computes a circular original-biarc-covering biwalk of $DBG'_k(I)$.*

*Proof.* We need to prove that given a solution to the flow formulation, the graph gets correctly biEulerised such that it admits a biEulerian circuit after step 4.

First, consider that due to Equation (2), the biEulerisation itself is well-defined, as there are only whole-numbered amounts of arcs to be added, and there are no negative numbers implying removal of arcs. Further, it holds that the biEulerised graph is a valid bigraph, since Equation (6) ensures that whenever an arc gets added, the corresponding reverse complement gets added as well.

Next, we show that Equations (4) and (5) ensure that the flow results in a valid biEulerisation, i.e. that the bi-imbalance of each node is zero after applying the biEulerisation. We define $x'_e$ be the multiplicity of each arc after applying the biEulerisation, defined as $x'_e := x_e + 1$ for non-breaking arcs and $x'_e := x_e$ for breaking arcs.

- If $v$ is not self-complemental, then it is constrained by Equation (4). We can rewrite it as follows:

$$\sum_{e \in E^-(v)} x'_e \left(1 + \mathbb{1}_{e=e^{-1}}\right) = \sum_{e \in E^+(v)} x'_e \left(1 + \mathbb{1}_{e=e^{-1}}\right). \tag{7}$$

  And, using $d'^+$ and $d'^-$ to denote degrees in the biEulerised graph as well as $\in_\#$ to denote the operator that counts the number of occurrences of an element in a multiset, we get:

$$0 = d'^+(v) - d'^-(v) + (((v, v^{-1}) \in_\# E^+(v)) - ((v^{-1}, v) \in_\# E^-(v))).$$

  The right part of the equation is the generalised formula for the bi-imbalance, allowing for duplicate self-complemental arcs. Since it is zero, it holds that the bi-imbalance of $v$ is zero.

- If $v$ is self-complemental, then it is constrained by Equation (5). We can rewrite it as follows:

$$\left(\sum_{e \in E^+(v)} x'_e\right) \bmod 2 = 0. \tag{8}$$

  Using $d'^+$ to denote the out-degree in the biEulerised graph, we get:

$$d'^+(v) \bmod 2 = 0. \tag{9}$$

  The right part of the equation is the formula for the bi-imbalace for self-complemental nodes. Since it is zero, it holds that the bi-imbalance of $v$ is zero.

Since the bi-imbalance of all nodes is zero, by definition of the bi-imbalance, the biEulerised graph admits a biEulerian circuit. This is computed and output in step 5. The result is a circular original-biarc-covering biwalk, because by Equation (3), the biEulerisation and therefore also the final graph contain a breaking arc. □

**Lemma 11** (Chinese postman costs equal flow costs)**.** *Let $DBG'_k(I)$ be the transformed graph of $I$. Let $x$ be a solution to the flow formulation of $DBG'_k(I)$. Let $w$ be a minimum-cost circular original-biarc-covering biwalk of $DBG'_k(I)$. If each non-breaking arc $e \in F$ appears in $x_e + 1$ biarcs in $w$, and each breaking arc $e \in (E \setminus F)$ appears in $x_e$ biarcs in $w$, then $c(w) = |F| + \sum_{e \in E,\ e \text{ is canonical}} c_e x_e$.*

*Proof.* The costs $c(e)$ of arcs $e$ in the circular original-biarc-covering biwalk are equal to the costs of arcs $c_e$ in the flow formulation. Additionally, for each biarc $[e, e^{-1}]$, by definition it holds that $c_e = c_{e^{-1}}$.

Let $e \in E$ be an arc and $e^{-1} \in E$ its reverse complement. Since only one of them is canonical (or they are equal), $e$ and $e^{-1}$ together contribute $c_e \cdot x_e$ in the flow formulation.

On the other hand, since $e$ appears in $x_e + 1$ (or $x_e$) biarcs in $w$, it holds that $e^{-1}$ appears in the same $x_e + 1$ (or $x_e$) biarcs in $w$. Therefore, together they contribute $(x_e + 1) \cdot c_e$ to the costs of $w$ if they are non-breaking, and $x_e \cdot c_e$ if they are breaking. If they are non-breaking, it holds that $c_e = 1$, and therefore $(x_e + 1) \cdot c_e = x_e \cdot c_e + 1$. Resulting, since there are $|F|$ non-breaking arcs, it holds that $c(w) = |F| + \sum_{e \in E,\ e \text{ is canonical}} c_e x_e$. □

**Lemma 12** (Min-cost flow can reproduce solutions at the same costs)**.** *Given a circular original-biarc-covering biwalk $w$ in a transformed graph $DBG'_k(I)$, there exists a solution $x$ to the flow formulation of $DBG'_k(I)$ that produces a circular original-biarc-covering biwalk $w'$ if fed into steps 4 and 5 of Algorithm 9, where the $c(w) = c(w')$.*

*Proof.* We construct $x$ by setting $x_e$ to the number of occurrences of arc $e$ in $w$, and then subtracting one from each $x_e$ if $e \in F$. Then, by Lemma 11, it holds that $c(w) - |F|$ equals the costs of $x$. Further, the circular original-biarc-covering biwalk $w'$ produced by 4 and 5 of Algorithm 9 contains $x_e$ occurrences of each arc $e \in E$, plus one additional occurrence of each non-breaking arc $e \in F$. Since each arc in $e \in F$ has costs $c_e = 1$, it holds that $c(w') - |F|$ equals the costs of $x$, so $c(w) = c(w')$.

It remains to show that $x$ is a valid solution to the flow formulation. We show that each condition holds.

(2) Holds.

(3) By definition, $w$ contains at least one breaking biarc, which by definition is an arc $e \in (E \setminus F)$. For this $e$ it holds that $x_e \geq 1$.

(4) Let $v \in V$ be a node that is not self-complemental. We can rewrite Equation (4) as follows:

$$\begin{aligned} &d_F^-(v) + \mathbb{1}_{(v^{-1},v) \in F^-(v)} + \sum_{e \in E^-(v)} x_e \left(1 + \mathbb{1}_{e=e^{-1}}\right) \\ &= d_F^+(v) + \mathbb{1}_{(v,v^{-1}) \in F^+(v)} + \sum_{e \in E^+(v)} x_e \left(1 + \mathbb{1}_{e=e^{-1}}\right) \end{aligned}$$

By defining $x'_e := x_e$ for breaking arcs and $x'_e := x_e + 1$ for non-breaking arcs, we can further rewrite the equation:

$$\sum_{e \in E^-(v)} x'_e \left(1 + \mathbb{1}_{e=e^{-1}}\right) = \sum_{e \in E^+(v)} x'_e \left(1 + \mathbb{1}_{e=e^{-1}}\right) \tag{10}$$

By definition, $x'_e$ is the number of occurrences of arc $e$ in $w$. We decompose $w = ([e_1, e_1^{-1}], \ldots, [e_{|w|}, e_{|w|}^{-1}])$ into two circular walks $w_1, w_2$ where $w_1 := (e_1, \ldots, e_{|w|})$ and $w_2 := (e_{|w|}^{-1}, \ldots, e_1^{-1})$. The number of occurrences of each arc in $w$ is the same as the sum of occurrences in $w_1$ and $w_2$, except for self-complemental arcs which occur twice as often in $w_1$ and $w_2$. Therefore, Equation (10) counts the sum of arcs in $w_1$ and $w_2$ that enter $v$ on the left side, and the sum of arcs that leave $v$ on the right side. Since $w_1$ and $w_2$ are circular walks, this implies that the sums are equal.

(5) Let $v \in V$ be a self-complemental node. By defining $x'_e := x_e$ for breaking arcs and $x'_e := x_e + 1$ for non-breaking arcs, we can rewrite Equation (5) as follows:

$$\left( \sum_{e \in E^+(v)} x'_e \right) \bmod 2 = 0 \tag{11}$$

By definition, $x'_e$ is the number of occurrences of arc $e$ in $w$. Since $v$ is self-complemental, it can only be entered or left via arcs that are not self-complemental. Therefore, whenever $w$ contains $v$, it contains one arc that leaves $v$, and another arc that leaves $v^{-1}$ in its reverse complement. Since $v = v^{-1}$, each element of the sum in Equation (11) is even, so the whole sum is even, so the equation holds.

(6) If $e \in E$ is self-complemental, this condition is a tautology. If not, then each biarc in $w$ that contains $e$ also contains $e^{-1}$, so it holds that $x_e = x_{e^{-1}}$. □

**Lemma 13.** *Algorithm 9 computes a minimum-cost circular original-biarc-covering biwalk $w$ of $DBG'_k(I)$.*

*Proof.* By Lemma 12 it holds that for a minimum-cost circular original-biarc-covering biwalk $w'$, there exists a flow solution that produces another (or the same) minimum-cost circular original-biarc-covering biwalk $w$. By Lemma 10, it holds that Algorithm 9 computes a circular original-biarc-covering biwalk of $DBG'_k(I)$, so it cannot compute any walk of lower costs, otherwise $w'$ would not have been minimum-cost. □

From Lemma 13 it follows that we can use the flow formulation with Algorithm 9 to solve step 3 of Algorithm 4 and get a complete algorithm to compute matchtigs.

#### Solving the min-cost integer flow formulation with min-cost matching.

As argued above, the flow formulation is not actually practically useful, therefore we solve it via min-cost perfect matching. For this, we are given the set of min-cost paths between unbalanced nodes as follows:

**Definition 14** (Matching paths)**.** *Let $DBG_k(I)$ be an arc-centric de Bruijn graph. Let $S$ be the set of its self-complemental nodes with positive bi-imbalance, $A$ be the set of its nodes with negative bi-imbalance and $B$ be the set of its nodes with positive bi-imbalance that are not self-complemental. We define $P := \{(u, v, c(u, v)) \mid u \in A \cup S \wedge v \in B \cup S \wedge c(u, v) \leq k - 1\}$ as the* set of min-cost source-sink paths *in $DBG_k(I)$, where $c(u, v)$ are the costs of the min-cost path from $u$ to $v$ where all arcs have costs of one. We additionally define $P_{u,v} := \min\{(\{u, v\}, c) \mid \exists u' \in \{u, u^{-1}\}, v' \in \{v, v^{-1}\} : (\{u', v'\}, c) \in P \vee (\{v', u'\}, c) \in P\}$ as the minimum min-cost path between $u$ and $v$ or their reverse complements, where min-cost paths are compared by their costs $c$. If there are no min-cost paths between $u$ and $v$ (i.e. the minimum is taken from the empty set), then $P_{u,v} := \bot$.*

We informally describe the matching instance in Section 5.6, so we formally restate it here for the purpose of our proof.

**Definition 15** (Matching instance)**.** *Let $DBG_k(I)$ be an arc-centric de Bruijn graph. Let $S$ be the set of its self-complemental nodes with positive bi-imbalance, $A$ be the set of its nodes with negative bi-imbalance and $B$ be the set of its nodes with positive bi-imbalance that are not self-complemental.*

*Then the matching graph is defined as:*

$$\begin{aligned} M &= (V, E) \\ V &= V_1 \cup V_2 \\ V_1 &= \{v_i+, v_i- \mid v \in (A \cup B \cup S) \wedge v \textit{ is canonical} \wedge i \in \{1, \dots, |bi_v|\}\} \\ V_2 &= \{u+, u-, w+, w-\} \text{ // extra nodes for Equation (3)} \\ E &= E_1 \cup E_2 \cup E_3 \end{aligned}$$

*where*

- $E_1 = \{(\{v_i+, v_i-\}, k-1) \mid v_i+ \in V_1\}$ *are the edges that allow nodes to stay "unmatched" (note that condition $v_i+ \in V_1$ is equivalent to $v_i- \in V_1$ by definition of $V_1$),*
- $E_2 = \{(\{v_i+, v'_j+\}, c), (\{v_i-, v'_j-\}, c) \mid (\{v, v'\}, c) = P_{v,v'} \wedge v_i+, v'_j+ \in V_1\}$ *are the edges for sending flow between $v$ and $v'$ to balance them, and*
- $E_3 = \bigcup_{v_i+ \in V_1} \{(\{u+, v_i+\}, 0), (\{w+, v_i+\}, 0), (\{u-, v_i-\}, 0), (\{w-, v_i-\}, 0)\}$ *are the edges allowing one pair of nodes to be connected via a breaking biarc for free.*

We use the matching instance to solve the flow formulation as follows:

**Algorithm 16.**

1. *Compute the arc-centric de Bruijn graph $DBG_k(I)$.*
2. *Compute the set of min-cost source-sink paths $P$ in $DBG_k(I)$.*
3. *Build and solve the matching instance $M = (V_1 \cup V_2, E_1 \cup E_2 \cup E_3)$ using a min-cost perfect matching algorithm.*
4. *Compute the transformed graph $DBG'_k(I) = (V, E)$.*

5. *Initialise the solution to the flow formulation with $x_e = 0$ for all $e \in E$.*

6. *For each arc $p \in E_2$ part of the matching solution, and for each min-cost path in $P$ represented by $p$, increment the flow values $x_e$ of all arcs $e \in E$ that are on a min-cost path by one.*

For showing that this algorithm computes a minimum-cost flow, we show that valid (not necessarily minimum) perfect matchings and (not necessarily minimum) flows are equivalent in terms of costs. But perfect matchings actually just represent a subset of possible flows. However, if a flow cannot directly be transformed into a matching, it is not minimum.

**Definition 17** (Path-minimum flow)**.** *A* path-minimum flow *is a not necessarily minimum flow in the flow formulation of a transformed graph $DBG'_k(I) = (V, E)$ such that the flow can be decomposed into a set of source-sink paths such that each path is a shortest path if each arc has weight one. A source is a node with negative imbalance or a self-complemental node with positive imbalance, and a sink is a node with positive imbalance.*

**Lemma 18.** *Each minimum flow is path-minimum.*

*Proof.* Let $x$ be a minimum flow in the flow formulation. Equations (4) and (5) ensure that the flow can be decomposed into a set of source-sink paths. If any of these paths were not min-cost, it could be replaced with a min-cost path, resulting in a cheaper flow and contradicting the minimality of $x$. Therefore, $x$ is path-minimum. □

**Lemma 19.** *Each path-minimum flow $x$ of costs $c(x)$ can be transformed into a perfect matching $E_M$ of costs $c(E_M) = 2(c(x) - (k-1))$.*

*Proof.* We construct the perfect matching $E_M$ as follows:

- Pick some breaking arc $e_b$ with positive flow (it exists because of Equation (3)) and add the edges $\{u+, v_1+\}, \{w+, v'_1+\}, \{u-, v_1-$[illegible] and $\{w-, v'_1-\}$ corresponding to $e_b$ to $E_M$.

- For each breaking arc $e$, repeat the following $x_e$ times, or $x_e - 1$ times if $e = e_b$: add the corresponding edges of the form $\{v_i+, v_i-\}$ to $E_M$, choosing an arbitrary $i$ such that no node gets matched twice.

- For each source-sink biwalk $p$ in the decomposed flow that contains no breaking biarcs, add the edges $\{v_i+, v'_j+\}$ and $\{v_i-, v'_j-\}$ corresponding to $p$ to $E_M$. Choose an arbitrary $i$ and $j$ such that no node gets matched twice. Repeat this step according to the minimum flow value on any arc in $p$.

We show that $E_M$ is a perfect matching. In the decomposition of $x$, each unbalanced binode is start or end of as many biwalks as there are copies of it in the matching instance. This specifically holds for walks from a binode to its complement binode, which alter the bi-imbalance of the corresponding binode by two, and also connect two nodes in the matching problem that correspond to the same binode. Since the matching instance has copies of each binode according to its absolute bi-imbalance, this means that all nodes in the matching instance are matched exactly once. Further, the arbitrarily picked arc $e_b$ ensures that also $u+, u-, w+$ and $w-$ are matched. So, $E_M$ is a perfect matching.

In $E_M$, pairs of edges correspond source-sink biwalks in $x$. In the flow formulation the costs of each biwalk are counted only in one direction, while in the matching instance, the costs are counted twice. Therefore, the costs of $E_M$ are twice those of $x$. However, the nodes corresponding to the picked arc $e_b$ are matched at no cost, whereas $x$ contains a breaking arc of costs $k-1$ between them. Therefore, the costs of $E_M$ are $c(E_M) = 2(c(x) - (k-1))$. □

**Lemma 20.** *Each perfect matching $E_M$ of costs $c(E_M)$ can be transformed into a path-minimum flow $x$ of costs $c(x) = c(E_M)/2 + (k-1)$.*

*Proof.* Note that each solution of the matching problem can be transformed into a solution of the matching problem which is symmetric, meaning that if an arc $\{v_i+, v'_j-\}$ is in $E_M$, then also $\{v_i-, v'_j+\}$ is in $E_M$, as well as if n arc $\{v_i+, v'_j+\}$ is in $E_M$, then also $\{v_i-, v'_j-\}$ is in $E_M$. Using this symmetry, we construct the path-minimum flow $x$ as follows:

- For each matched edge of the form $\{v_i+, v'_j+\}$ and $\{v_i-, v'_j-\}$, we add a flow of one to the corresponding biwalk in $x$.

- For each pair of distinct matched edges of the form $\{v_i+, v_i-\}$, $\{v'_j+, v'_j-\}$ we add flow to the corresponding breaking biarc in $x$.

- For the matched edges $\{u+, v_1+\}, \{w+, v'_1+\}, \{u-, v_1-\}$ and $\{w-, v'_1-\}$ we add flow to the corresponding breaking biarc in $x$.

We show that $x$ is a valid flow. Each unbalanced binode has copies in the matching instance according to its absolute bi-imbalance. Therefore, in the constructed flow $x$, source and sink nodes are balanced. Also, all other nodes are balanced, as they were balanced before, and are only passed via paths that keep them balanced. Therefore, $x$ is a valid flow, and by construction it is path-minimum.

In $E_M$, pairs of edges correspond to source-sink biwalks in $x$. In $x$ the costs of each biwalk are counted only in one direction, while in the matching instance, the costs are counted twice. Therefore, the costs of $E_M$ are twice those of $x$. However, the nodes corresponding to the picked arc $e_b$ are matched at no cost, whereas $x$ contains a breaking arc of costs $k-1$ between them. Therefore, the costs of $x$ are $c(x) = c(x)/2 + (k-1)$. □

**Theorem 21.** *The matchtigs algorithm composed of Algorithms 4, 9 and 16 is correct.*

*Proof.* Algorithm 4 is correct by Lemma 7. Algorithm 9 is correct by Lemma 13. Algorithm 16 computes a min-cost flow because minimum-cost flows are path-minimum by Lemma 18, and path-minimum flows are equivalent to perfect matchings in terms of costs by Lemmas 19 and 20. □
