## Additional file 10 for "Matchtigs: minimum plain text representation of kmer sets"

### Additional file 10: query experiment on a machine with focus on single-core performance

Table 1: Performance characteristics of querying different tigs with SSHash-Lite.

(a) regular SSHash-Lite

| genome | algorithm | index time [min] | search time [sec] | search speedup | | index size [GiB] | size imprv. | |
|---|---|---|---|---|---|---|---|---|
| ~309kx | unitigs | 1.86 | 1084 | 1.00 | | 0.99 | 1.00 | |
| Salmonella | UST | 1.38 | 568 | 1.91 | | 0.67 | 1.48 | |
| (0.75) | gMatchtigs | 1.48 | 339 | 3.20 | (1.68) | 0.62 | 1.60 | (1.08) |
| Human | unitigs | 13.8 | 274 | 1.00 | | 4.39 | 1.00 | |
| reads | UST | 11.4 | 252 | 1.10 | | 3.46 | 1.27 | |
| (0.75) | gMatchtigs | 12.9 | 208 | 1.31 | (1.2) | 3.31 | 1.33 | (1.05) |
| 2505x | unitigs | 9.50 | 249 | 1.00 | | 3.57 | 1.00 | |
| Human | ProphAsm | 8.58 | 215 | 1.16 | | 2.84 | 1.26 | |
| (0.65) | gMatchtigs | 9.02 | 188 | 1.32 | (1.14) | 2.84 | 1.26 | (1.00) |

(b) canonical SSHash-Lite

| genome | algorithm | index time [min] | search time [sec] | search speedup | | index size [GiB] | size imprv. | |
|---|---|---|---|---|---|---|---|---|
| ~309kx | unitigs | 2.81 | 413 | 1.00 | | 1.08 | 1.00 | |
| Salmonella | UST | 2.11 | 286 | 1.44 | | 0.75 | 1.44 | |
| (0.75) | gMatchtigs | 2.36 | 211 | 1.96 | (1.36) | 0.71 | 1.52 | (1.06) |
| Human | unitigs | 19.5 | 156 | 1.00 | | 4.79 | 1.00 | |
| reads | UST | 16.3 | 145 | 1.08 | | 3.86 | 1.24 | |
| (0.75) | gMatchtigs | 17.7 | 129 | 1.18 | (1.09) | 3.76 | 1.27 | (1.03) |
| 2505x | unitigs | 12.0 | 134 | 1.00 | | 3.93 | 1.00 | |
| Human | ProphAsm | 13.1 | 122 | 1.10 | | 3.19 | 1.23 | |
| (0.65) | gMatchtigs | 13.2 | 114 | 1.18 | (1.07) | 3.22 | 1.22 | (0.99) |

SSHash-Lite is run with $k = 31$ and a kmer-inclusion rate of 0.8. On the Salmonella pan-genome we used a minimizer length of 17 for the regular index and a minimizer length of 16 for the canonical index. On the human reads we used a minimizer length of 20 for the regular index and a minimizer length of 19 for the canonical index. On the human pangenome we used a minimizer length of 19 for the regular index and a minimizer length of 20 for the canonical index. The search speedup is with respect to unitigs, and the search speedup in parentheses is with respect to the strings computed by UST. Index time is the end-to-end time required to build the SSHash-Lite index: it includes reading the collections from disk and building the data structure using external memory. Searching time is the time required to check which reads have at least 80% of their kmers in the input SPSS. The number in parentheses under the genome is the kmer hitrate, i.e. the fraction of kmers from the query that are part of the queried dataset. These experiments were performed on an Intel Core i9-9900K CPU, clocked at 3.60 GHz and the code was compiled with gcc 11.2.
